# Attenuation of epigenetic regulator SMARCA4 and ERK-ETS signaling suppresses aging-related dopaminergic degeneration

**DOI:** 10.1101/835876

**Authors:** Ling Sun, Jie Zhang, Wenfeng Chen, Yun Chen, Xiaohui Zhang, Mingjuan Yang, Min Xiao, Fujun Ma, Yizhou Yao, Meina Ye, Zhenkun Zhang, Kai Chen, Fei Chen, Yujun Ren, Shiwei Ni, Xi Zhang, Zhangming Yan, Zhi-Rong Sun, Hai-Meng Zhou, Hongqin Yang, Shusen Xie, M Emdadul Haque, Kun Huang, Yufeng Yang

## Abstract

Parkinson’s disease (PD) is a complex disease with high heterogeneity. How complex interactions of genetic, environmental factors and aging jointly contribute to dopaminergic degeneration in PD is largely unclear. Here, we applied frequent gene co-expression analysis on human patient substantia nigra-specific microarray datasets to identify potential novel disease-related genes. *In vivo Drosophila* studies validated two of 32 candidate genes, a chromatin remodeling factor SMARCA4 and a biliverdin reductase BLVRA. Inhibition of SMARCA4 was able to prevent dopaminergic degeneration not only caused by overexpression of BLVRA but also in four most common *Drosophila* PD models. Mechanistically, aberrant SMARCA4 and BLVRA converged on elevated ERK-ETS activity, attenuation of which by either genetic or pharmacological manipulation effectively suppressed dopaminergic degeneration *in vivo*. Drug inhibition of MEK/ERK also mitigated mitochondrial defects in PD gene-deficient human cells. Our findings underscore the important role of epigenetic regulators and implicate a common signaling axis for therapeutic intervention in a broad range of aging-related disorders including PD.

## BACKGROUND

Among aging-related diseases, Parkinson’s disease (PD) is the most common neurodegenerative movement disorder, with an incidence rate above 1% among individuals over 65 years of age [1]. The pathologic manifestations of PD include age-dependent progressive dopaminergic (DA) neuronal deterioration in basal ganglia and *substantia nigra,* with reduction of dopamine release. Remarkable similarities at the molecular and cellular levels exist between PD and normal aging. Current treatments for PD are only symptomatic, ameliorating disease symptoms for a limited period of time, without retarding or halting disease progression.

PD is a complex disease with high heterogeneity. Etiologically, PD consists of early-onset subtypes, which are primarily due to high penetrance mutations and familial inheritance, and late-onset subtypes, which occur more sporadically and are believed to result from complex interactions between genetic, environmental factors superimposed on the physiological decline of neuronal functions with age. Emerging evidence affirms the central role of genetic susceptibility in PD [1, 2]. Although a comprehensive genetic architecture corresponding to distinct PD subtypes remains poorly understood, the same set of susceptibility genes may predispose people to both familial- and sporadic-PD. A number of causal genetic risk factors have been linked to PD onset, including mutations in *SNCA* (α-synuclein), *LRRK2* (leucine-rich repeat kinase 2), *VPS35* (the vacuolar sorting protein 35 gene), *EIF4G1* (eukaryotic translation initiation factor 4-gamma) and *DNAJC13* [DnaJ heat shock protein family (Hsp40) member C13] genes with autosomal dominant inheritance mode, and *PARK2* (parkin), *PINK*1 (PTEN induced putative kinase 1), *PARK7* (Parkinsonism associated deglycase, *DJ1*), *HTRA*2 (high temperature requirement A2), *DNAJC*6 [DnaJ heat shock protein family (Hsp40) member C6], *FBXO7* (F-box domain-containing protein), *PLA2G6* (phospholipase A2 group VI), *SYNJ1* (synaptojanin 1), *ATP6AP2* (ATPase H^+^ transporting accessory protein 2) and *ATP13A2* (ATPase type 13A2) with recessive inheritance mode [3].

Typically, these known PD genes participate in diverse cellular processes; however, common themes in PD pathogenesis have been proposed, such as aberrant proteostasis and vesicle trafficking, mitochondrial dysfunction, altered epigenetic regulation and inflammation [4]. Notably, all these involved pathogenic agents are similar to those in normal aging. Therefore, the insights involved in PD pathogenesis may be critical for understanding and modifying aging, and vice versa. Nevertheless, the majority of PD heritable components remain elusive [1, 2]. How to extract contributing factors from limited human brain specimens is the main challenge to solve this heterogeneous disease, because only postmortem human brain specimens can be available. More challengingly, most postulated novel genetic associations or risk factors await further validation.

Gene co-expression analysis allows identifying genes with similar expression patterns across a set of samples, which has facilitated identifying genes involved in certain disease pathways, new gene functions, and potential biomarkers [5, 6]. In this study, we used the known PD genes as “anchors” in order to identify new PD candidate genes that are highly co-expressed with known PD genes with multiple human brain microarray datasets. Among the predicted 32 candidate genes, SWI/SNF related, matrix associated, actin dependent regulator of chromatin, subfamily A, member 4 (*SMARCA*4) and biliverdin reductase A (*BLVRA*) were further studied *in vivo* using *Drosophila melanogaster*, and their potential involvement in PD pathogenesis were confirmed. Furthermore, we revealed a potential common aging-related pathogenic signaling pathway consisting of the chromatin-remodeling factor SMARCA4 and the ERK-ETS signaling axis, suggesting new therapeutic targets for PD and other ageing-related disorders, as three ERK-ETS inhibitors were tested for their efficacy in multiple *Drosophila* PD models. Our work also illustrates the high efficiency of combining bioinformatics analysis of large-scale human transcriptomic data and small-scale genetic screening using model organisms to interrogate highly heterogeneous diseases including aging-related disorders.

## METHODS

### Frequent gene co-expression analysis

We chose seven of the most commonly known PD genes as anchor genes. They are ATP13A2, HTR2A, SCNA, LRRK2, PARK2, PARK7, and PINK1. In addition, we identified eleven gene expression datasets from NCBI Gene Expression Omnibus (GEO), which contain samples from human brain tissues, especially the *substantia nigra* region in which the death of dopamine contain cells leads to PD. They are: GDS2519, GDS2821, GDS3128, GDS3129, GSE19587, GSE20141, GSE20146, GSE20153, GSE20292, GSE20295, and GSE20333.

Our workflow is similar to previously described in Conference papers, with a slight modification as in the following steps:

Step 1: For the i-th dataset (i= 1, 2, …, 11), compute the Pearson correlation coefficients (PCC) between every pair of genes within each dataset, and set the top five percentile of all PCC values as threshold Ti. PCC values were converted to absolute values before setting the threshold.
Step 2: For the k-th anchor gene Ak (k = 1, …, 7), denote 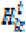 as the set of genes in the i-th dataset whose PCC values with respect to Ak are higher than Ti. These genes are considered to have high correlation with Ak in the i-th dataset.
Step 3: For a gene Gj, its frequency of having high correlation with Ak is denoted as

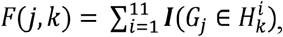

Where 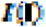 is the indicator function which is 1 if the input is TRUE and 0 otherwise.
Step 4: For each anchor gene Ak, let Pk be the set of genes with high 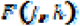 values. Specifically,

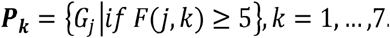

which is the collection of all genes that have high PCC values with Ak in at least five datasets for all seven anchor genes.
Step 5: Finally, for every gene Gj, count the frequency that it appears in Pk (k =1, …, 7). The count number of each gene Gj can be derived as

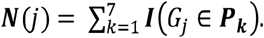

Finally, the selected gene networks are subjected to gene set enrichment analysis using TOPPGene (http://toppgene.cchmc.org/) and Ingenuity Pathway Analysis (IPA®, http://www.ingenuity.com).

The 32 PD candidate gene list is the result of a very stringent selection process, which involves three thresholds for gene selection. First, in each dataset only genes with top 5-percentile correlation coefficients for a specific anchor gene is selected for the next step. Second, only genes who have strong correlation (within top 5-percentile) with at least three anchor genes (out of total of seven anchor genes) are further selected as the gene list for this dataset. Third, genes that are selected in at least five datasets out of a total of 11 datasets are used in this study. Statistical analysis (based on Fisher’s exact test) shows that the chance for a random gene to be selected through this process is 3.4E-10.

### *Drosophila* Stocks and nomenclature

Fly strains were maintained with standard molasses-cornmeal-yeast food and were kept in 25°Cwhereas PD model flies were maintained at 21-25°C routinely and raised at 29°C for the experiments unless mentioned. TH-Gal4 was a gift from S. Birman [7]. Brm∷GFP is the Bloomington stock #59784. UAS-dBVR plasmid was derived from fly cDNA and molecular cloning, and the corresponding p-element mediated transgenic flies were generated by standard microinjection protocol using *w*^*1118*^ stock. Two UAS-dBVR transgenic lines were used in this study, UAS-dBvr and UAS-dBvr (III). Pnt∷EGFP is the Bloomington stock #42680 or #60528.

UAS-Brm wt, UAS-Brm^DN^, UAS-Brm RNAi fly strains were gifts from Helena E. Richardson. UAS-Lrrk2^I1915T^ and UAS-Pink1 RNAi fly strains were gifts from Bingwei Lu [8]. UAS-αSynA30P was a gift from Nancy Bonini [9]. The *pink1*^*B9*^ fly strain was a gift from Jongkyeong Chung. Parkin RNAi is the Bloomington stock #38333. GstD-GFP was a gift from Dirk Bohmann. Other fly strains were obtained from Bloomington *Drosophila* Stock Center (#31603, #33639, #34855, #35038, #869, #5789, #5790, #34909, #59006, #67672) and Vienna *Drosophila* Research Center (dBVR RNAi, v24042). The age of adult flies was defined as the days after eclosion (AE). Further information on genes and symbols can be found in Flybase (http://flybase.bio.indiana.edu). Fly genotypes for each experiment were listed in the supplementary appendix.

### SNP query method

We queried with gene symbol “SMARCA4” and “BLVRA” in PD gene database PDGene (http://www.pdgene.org), which described that the SNP data source as “the data” currently available on PDGene include all results pertaining to the discovery phase of the GWAS meta-analysis by [1]. This includes data on 7,782,514 genetic variants in up to 13,708 PD cases and 95,282 controls from 15 independent GWAS datasets of European descent. Variants were imputed using the August 2010 release of the 1000 Genomes Project European-ancestry haplotype reference set and filtered according to standard quality control criteria. In line with the criteria applied in the published study [1], only variants with a minor allele frequency ≥0.1% and those assessed in at least 3 of the 15 datasets have been included for display in PDGene. The database also includes association results of genotyping data generated on the “NeuroX chip”, a semi-custom genotyping array, on 5,353 PD cases and 5,551 controls of European descent for the most significantly associated polymorphisms from the discovery phase (i.e. across 26 loci showing genome-wide significant association (p <5×10^−8^) in the discovery phase with PD risk) as well as for 6 additional, previously reported GWAS signals. Details on the included datasets as well as all genotyping procedures and statistical analyses can be found in our original publication [1].

### *Drosophila* PD models and pathologic phenotype evaluation

The following genotype-based *Drosophila* PD models were used in this study: 1) TH-Gal4>UAS-αSynA30P; 2) TH-Gal4>UAS-Lrrk2^I1915T^; 3) TH-Gal4>UAS-Parkin RNAi (Bloomington #38333); 4) *pink1*^B9^; TH-Gal4 or TH-Gal4>UAS-PINK1 RNAi. These four *Drosophila* PD models were abbreviated as αSyn, Lrrk2, Parkin, Pink1 (Pink1 mut and Pink1 RNAi) PD models respectively. Control flies were TH-Gal4>w- or TH-Gal4>UAS-Luc RNAi. Homozygous *parkin* null alleles were found not healthy, so they were not used for PD modeling in this study. Parkin RNAi flies were used instead. *pink1*^B9^ was primarily used as the *pink1*-related PD fly model, see genotype listed in the supplementary appendix. Male flies were used unless mentioned. Experimental flies were sorted into individual vials at a density of 15~20 flies per vial and were transferred to fresh vials three times a week. Experimental flies were raised at 29°C. DA neurons were marked by anti-tyrosine hydroxylase (TH) antibody. The DA neuron number in the lateral protocerebral posterior 1 (PPL1) cluster was scored. Left and right PPL1 clusters in individual fly brains were scored independently. Only well-dissected, processed, mounted and preserved fly brains were used for the quantifications. The intact left or right PPL clusters contains 11~12 DA neurons in healthy individuals. 2-day-old (or 2^nd^ day AE) and 30-day-old (or 30^th^ day AE) adult flies were subjected to evaluation of DA neurons for most experiments unless noted. At least 20 hemisphere brains were quantified via double-blinded fashion for each data point (n>20). Two biological replicates were carried out.

### Semi-quantitative RT-PCR and Quantitative Real-time PCR

Total RNA extracted from fly heads with RNA extraction kit (TRIZOL reagent, Invitrogen, Inc.) was used for semi-quantitative reverse transcription-PCR (Sq-RT PCR) and Quantitative real-time PCR (Q-RT PCR). The cDNA was synthesized using the Reverse Aid First strand cDNA synthesis kit (Thermo Fisher, Cat NO: K1622). Specific primer pairs were used for BVR and αSYN gene expression analysis. Expression levels of any given genes were resolved by agarose gel electrophoresis for the Semi-quantitative RT-PCR. The samples were analysis by Quantity One BioSoft (BIO-RAD). Relative expression levels were normalized to that of tubulin. Two biological replications were carried out for each sample. The Q-RT PCR was performed using a LightCycler® 96 System instrument with Ultra SYBR Mixture (CW Bio, CW0957M). The data display fold change relative to the control after normalization to a-Tubulin. More than three replications were carried out.

### Whole-mount Brain Immunostaining, live imaging and Microscopy

Adult flies were collected for brain dissection at the indicated time points. Brains were fixed for 2 hours in 4% buffered formaldehyde at 25°C, washed in phosphate-buffered saline (pH 7.4) with 0.2% Triton X-100 (PBT), blocked in 5% goat serum in PBT (PBST) for 30min at room temperature and incubated in primary antibody overnight at 4°C. Primary antibodies were prepared in blocking buffer solution (Rabbit anti-TH, Millipore, #AB152). After three times of 15-min-wash, brains were incubated with goat anti-rabbit secondary antibodies (Life AlexaFluo® 488, 568) for 2 h at room temperature, followed by thorough rinses and mounted. For live imaging, fly brains or imaginal discs were promptly dissected in *Schneider*’s insect medium (Life-Gibco), properly mounted and scanned, stained with Hoechst33342 (Beyotime) in some cases prior to imaging in some cases by standard protocol. Average fluorescence intensity of Brm∷GFP, Pnt∷EGFP or GstD-GFP was quantified with Metamorph software (Leica AF lite), normalized to those in control flies. At least five well-preserved fly brains were used for fluorescence intensity quantifications. All images were taken by a confocal microscopy (Leica TCS SP5) with identical instrument parameters for any given individual experimental series. Images were processed with Adobe Photoshop and subjected to identical post-acquisition brightness/contrast effects.

### Western Blot Analysis

Fly heads were homogenized with a pestle, and protein extracts were prepared with lysis buffer (20 mM Tris/HCl at pH 7.6, 150 mM NaCl, 5mM EDTA, 10% glycerol, 1% SDS and 1 mM PMSF). Supernatants were collected after 16000g and 4°C for 10 min, added with 5×SDS loading buffer and boiled for 5 min at 95°C. For immune-blotting analysis, protein lysates were electrophoresed with SDS-PAGE and transferred to PVDF membranes (Bio-Rad). Membranes were blocked in 5% BSA in PBS-T, followed incubation with diluted antibodies and secondary antibodies in blocking solution. Primary antibodies used were: monoclonal mouse anti-alpha tubulin (DSHB), monoclonal mouse anti-pERK (Sigma #M8159), rabbit anti-ERK (Cell Signalling #4695). Proteins were visualized using the Immobilon Western chemiluminescent HRP substrate (Beyotime) on ChemiDoc^TM^ XRS+ (BIO-RAD). Band intensity was calculated and analyzed with the Quantity One v4.62 (BIO-RAD). At least three biological replicates were performed and their means were calculated. Statistics were analyzed with Student *t*-test for numerical data.

HeLa cell lines were grown in Dulbecco’s Modified Eagle Medium (DMEM) containing 10% fetal bovine serum in a 5% carbon dioxide (CO_2_) atmosphere. Control and *pink1*-KO HeLa cells were homogenized with RIPA Lysis Buffer (Beyotime). Blots were probed with the following antibodies: monoclonal mouse anti-alpha tubulin (DSHB), rabbit anti-PINK1 (Cell Signalling #6946), monoclonal mouse anti-pERK (Sigma #M8159), rabbit anti-ERK (Cell Signalling #4695), rabbit anti-pMEK (Cell Signalling #3958), rabbit anti-MEK (Cell Signalling #13033).

### Drug treatment experiment

The experimental flies were collected after eclosion, assorted into 20 flies per via and raised at 29°C for drug treatment. U0126 [1,4-Diamino-2,3-dicyano-1,4-bis (o-aminophenylmercapto) butadiene, Selleck #S1102] were dissolved with DMSO in the recommended stock solution (10 mg/mL). Flies were fed for 4h with a serial concentration gradient of U0126, 10 μg/mL and 1 μg/mL, which were diluted with 4% sugar water, and were transferred back to standard fly food after drug exposure. Prior to drug treatment, a food dye supplement was used to justify the feasibility of ingestion of drugs by flies via this protocol. Drug treatment was performed continuously in a 24h cycle until the flies were harvested for protein analysis (7 days drug treatment) or whole-mount immunostaining analysis (30 days drug treatments). For control treatments, equivalent volumes of the vehicle alone were added. The application of the PD0325901 (Selleck #S1036) or Trametinib (Selleck #S2673) followed the identical procedure as U0126, with 1 μg/mL and 10 μg/mL feeding concentration for PD0325901, and 1.624 μM and 16.24 μM for Trametinib, respectivefully. Two independent sets of biological experiments were performed.

### Generation of *pink1* knockout HeLa cell lines using CRISPR/Cas9 gene editing

The HeLa cell line was sent to GENEWIZ, Inc. (Beijing, China) to perform authentication test. Firstly, genomic DNA was extracted from the cell pellets. Samples together with positive and negative control were amplified using GenePrint 10 System (Promega). Then, the amplified products were processed using the ABI3730xl Genetic Analyzer. Finally, data were analyzed using GeneMapper software V.4.0 and then compared with the ATCC for reference matching. To generate *pink1* knockout cell lines, CRISPR guide RNAs (gRNAs) were chosen to target exon 1 which is common to all splicing variants. Oligo nucleotides containing CRISPR target sequences (5’-CCGGCCGGGCCTACGGCTTG-3’) were annealed and ligated into pSpCas9 (BB)-2A-GFP (PX458) (Addgene 48138). Then, HeLa cells were transfected with this Cas9-2A-GFP and gRNA constructs. Two days after transfection, DNA from polled cells were extracted and the targeted genomic regions were PCR amplified. PCR products were subjected to Sanger sequencing analysis to verify the potential success of targeting. GFP-positive cells were sorted by FACS and plated in 96-well plates. Single colonies were expanded for depletion screening of the mutations. Knockout lines were further confirmed by Sanger sequencing. A cell clone harboring two heterogeneous frame-shift mutations at the *pink1* locus was used for subsequent experiments, referred as *pink1^−/−^*. Western blot analysis was conducted to validate the loss of Pink1 with the anti-PINK1 (D8G3) Rabbit mAb (Cell Signaling, #6946). DMSO was the solvent and equivalent amount was used in parallel as the drug treatment control.

### Assessment of mitochondrial membrane potential (MMP)

Mitochondrial membrane potential was assessed in WT and *pink1*-KO HeLa cells with the probe JC-1 (Invitrogen). JC-1 accumulates within the intact mitochondria to form multimer J-aggregates that result in a shift of fluorescence from green (530 nm) to red (590 nm). The potential-sensitive color shift is due to concentration-dependent formation of red fluorescent J-aggregates. A change of fluorescence from red to green indicates decreased MMP. Cells were treated with PD0325901 (50 nM), a selective and non ATP-competitive MEK inhibitor, for 8 hours. Then, the cells were loaded with 5 μg/ml of JC-1 for 3 minutes at 37°C. The cells were rinsed with phosphate-buffered saline, and mitochondrial JC-1 was analyzed by a Leica SP5 confocal microscopy and the Leica MetaMorph software. More than 30 randomly selected individual cells were analyzed for each data points. Three biological replicates were performed.

### Assessment of mitochondrial content and morphology

Mitochondrial content and morphology in WT and *pink1*-KO HeLa cells were visualized with Mito-Tracker Red (Molecular Probes) by a Leica SP5 confocal microscopy. Cells were treated with PD0325901 (50 nM) for 8 hours and then stained with MitoTracker for 30 min. Finally, the mitochondrial content and morphology were assessed with Mito-Morphology Macro in ImageJ as previously described [10, 11]. DMSO was the solvent and equivalent amount was used in parallel as the drug treatment control. More than 30 randomly selected individual cells were analyzed for each data points. Three biological replicates were performed.

### Assessment of whole brain Redox state

The CM-H2 DCFDA fluorescein dye (Invitrogen, Cat# C400) and redox-sensitive GFPs (roGFPs) protein were employed to measure the whole brain ROS stress of PD *Drosophila*. Measurement methods involved in were introduced from previously reported publications [12, 13]. The *Drosophila* brains with different genetic backgrounds were live dissected and incubated with 10 μM DCFDA for 5 min at RT if applicable, brain images were captured using a Leica TCS SP5 II confocal microscope with 488 nm excitation and 525 nm emission. Images were analyzed by image J software.

Alternatively, fly lines of tub-mito-roGFP2 or UAS-roGFP2 genotype were crossed with the PD models respectively in order to estimate the ROS level in whole brain or in the PPL1 neurons. The aged *Drosophila* brains were dissected in PBS with 20 mM NEM (N-ethyl maleimide) and then were imaged with a 535nm filter, followed with excitations at 405 nm and 488 nm. Image J software were employed to analyzed the 405 nm:488 nm ratios.

### Quantification and statistical analysis

Error bars represent standard deviations (S.D.) as indicated in the figure legends. Statistical analyses were performed using GraphPad Prism (GraphPad Software). Statistical analysis of differences between two groups was performed using Mann-whitney test. * indicates P < 0.05; **, P < 0.01; ***, P < 0.001; ns, not significant. Differences in means were considered statistically significant at P < 0.05.

## RESUTLS

### Gene co-expression network analysis identified 32 novel PD-associated candidate genes

We chose seven of the most commonly known PD genes as anchor genes, namely *SCNA, LRRK2, PARKIN, DJ1*, *PINK1, ATP13A2,* and *HTR2A.* Eleven datasets from NCBI Gene Expression Omnibus (GEO) were used, which contained samples from human brain tissues, especially the *substantia nigra* region. Our workflow is illustrated in Figure 1a. A total of 32 genes were identified to have high Pearson correlation coefficients (PCC) with at least three anchor genes in at least five datasets (Table 1 and Additional file 1: Table S1). According to the gene ontology (GO) enrichment analysis, the identified candidate PD genes were highly enriched with the genes associated with age-dependent metabolic reprogramming and neural disorders [14].

**Fig. 1.**
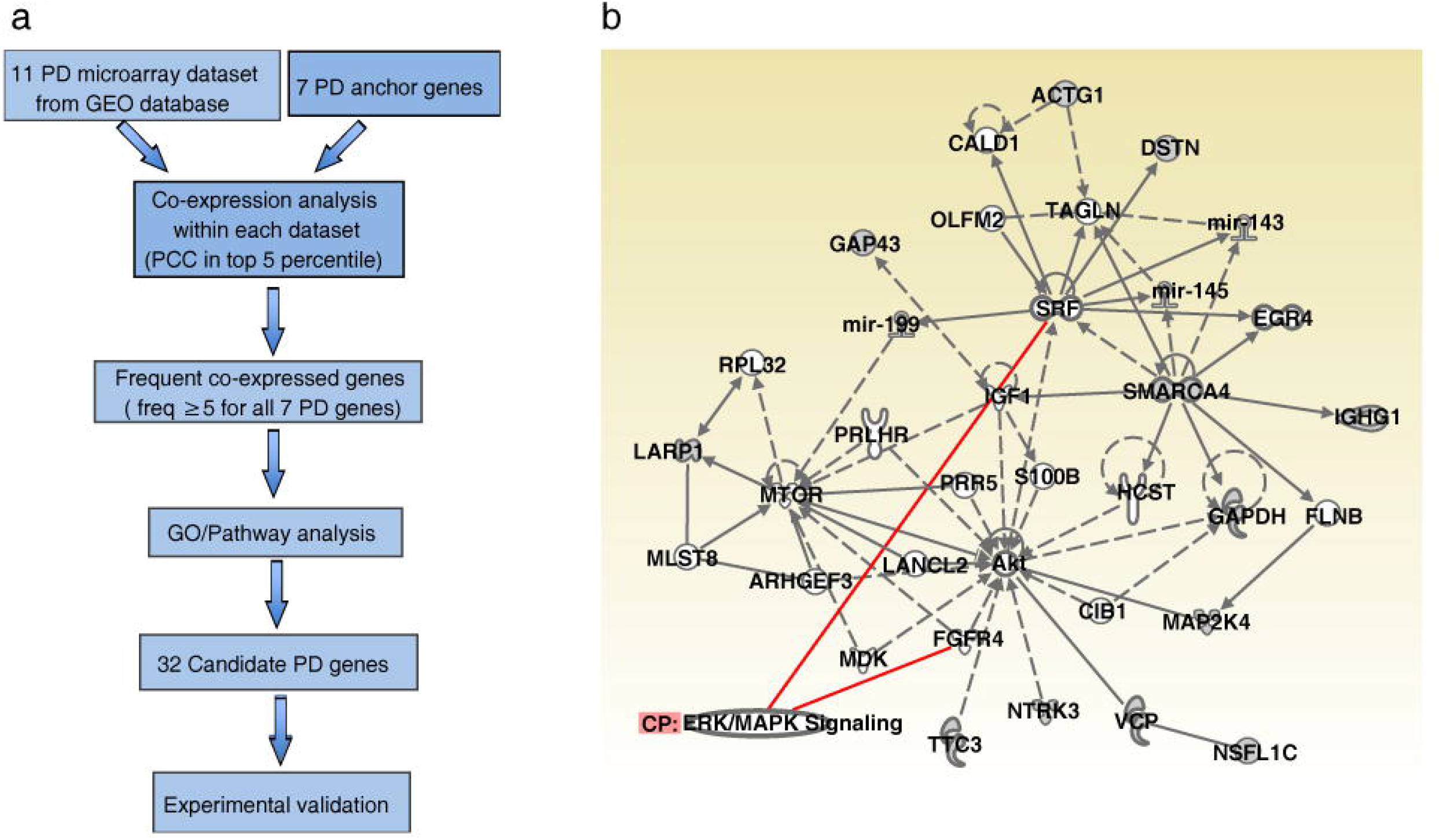
Bioinformatics analyses. **a** Overview of frequent gene co-expression analysis in identifying novel Parkinson’s disease-associated genes. Eleven NCBI GEO DataSets were identified with the microarray datasets of the substantia nigra brain region of human PD patients. Seven genes were used as the anchor PD genes. See Materials and Methods for more details. **b** Ingenuity Pathway Analysis (IPA) analysis revealed the connection of SMARCA4/Brm with MAPK kinase pathway. Red lines indicate genes known to interact with ERK signaling. Solid line: direct interaction; dash line: indirect interaction; arrows indicate the direction of interaction between two molecules. Different shapes indicate the type of the molecules.

**Table 1.** Genes with highest anchor gene count (*N*[·]values) in at least five datasets.

| Anchor Gene Count ($N\cdot$) | Gene Symbols |
| --- | --- |
| **5** | ***SMARCA4*** |
| 4 | *UBE3A, SNRPN, SLC25A3, PRDX2, GNAS, ARF1, ACTG1* |
| 3 | *VCP, TTC3, STMN2, RNF187, OBSL1, OAZ1, NTRK3, NSFL1C, MAP2K4, LARP1, KLC1, IGHG1, GAPDH, GAP43, EPB41L1, DSTN, DKK3, CLTA, BLVRA, ATP6V1C1, ATP5O, ATP5G3, ARF3, AAK1* |

### Inhibition of Brahma rescued DA degeneration caused by overexpression of BVR in *Drosophila*

Among the 32 candidate genes, we chose SMARCA4 (Brahma or Brm for *Drosophila* homologue) and BLVRA (biliverdin reductase A, dBVR for *Drosophila* homologue, CG9471) for further studies based on the rationales below (Additional file 1: Table S2). First, mutations of SMARCA4/Brm, a subunit of the SWI/SNF chromatin-remodeling complex that regulates higher order chromatin structure and gene expression, have been linked to multiple neurological and psychiatric disorders including autism spectrum disorders and schizophrenia [15, 16]. Although expression of SMARCA4/Brm has been reported in both murine and human DA neurons by recent single cell RNA-seq profiling [17, 18], there has been no functional reports of its role for DA neurons yet. Biliverdin reductases (BLVRs), together with hemeoxygenases (HOs), constitute the evolutionarily conserved enzymes in the heme metabolism, exert multiple physiological functions and have been considered as a potential biomarker for AD and mild cognitive impairment [19]. We then queried with gene symbol “SMARCA4” and “BLVRA” in PD gene database PDGene (http://www.pdgene.org), which incorporates all available SNP data pertaining to the discovery phase of the GWAS-meta-analysis [1]. We found that both SMARCA4 (Additional file 1: Table S3) and BLVRA (Additional file 1: Table S4) harbor SNPs with meta-analysis *P* value between 1E-4 and 0.05, which can be regarded as the potential PD risk SNPs albeit not in the top 10,000 most significant GWAS results.

To study the *in vivo* roles of candidate genes in PD pathogenesis, we took advantage of the *Drosophila melanogaster* model organism. The age-dependent progressive DA neuronal loss in the lateral protocerebral posterior 1 (PPL1) cluster was used as the neurodegenerative index (Fig. 2a-b). We then used an available Brm∷GFP reporter fly strain to examine whether Brahma is expressed in fly DA neurons [20]. Brm∷GFP was first verified by its nuclear localization in *Drosophila* larval tissues (Additional file 1: Figure S1a-c). Whole-mount immunostaining then confirmed the expression of Brm in the fly DA neurons (Fig. 2c), consistent with a previous report [21].

**Fig. 2.**
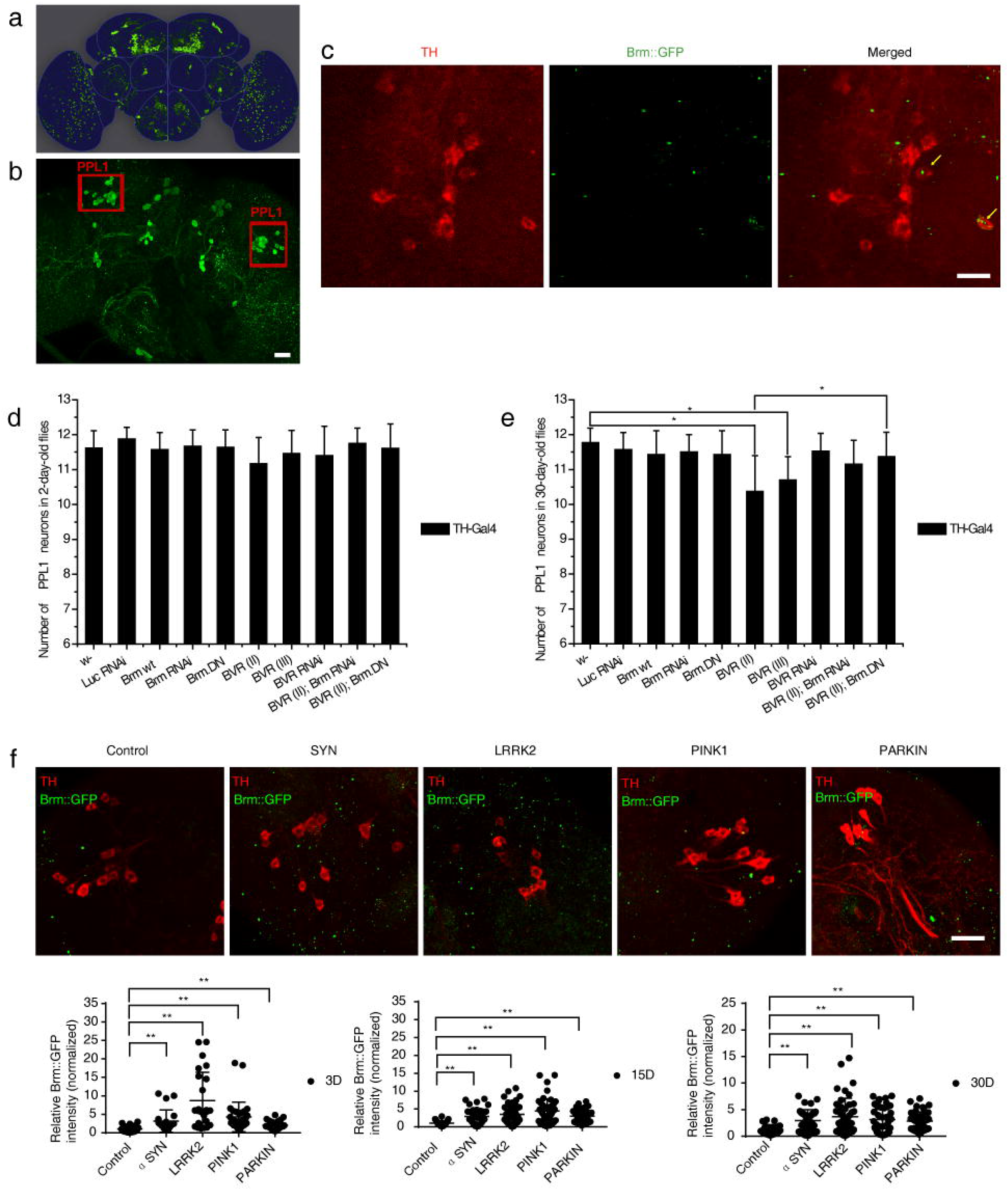
Inhibition of Brahma rescued dopaminergic (DA) degeneration caused by overexpression of BVR in *Drosophila.* **a** Schematic diagram of representative DA neuronal clusters in adult *Drosophila* brain. **b** Representative images show the posterior DA clusters in the fly brain. The intact left or right PPL cluster contains ~12 DA neurons at average in healthy adult flies. The DA neurons were labeled with green fluorescent protein (GFP). Scale bars, 25 μm. **c** Expression of Brm in the *Drosophila* DA neurons. A fusion Brm∷GFP transgene was used to report the endogenous protein expression level of Brm. The Brm protein (Green) was present in the *Drosophila* PPL1 DA neurons (anti-TH, Red) (pointed with yellow arrows) but not limited in DA neurons. Scale bars, 10 μm. **d, e** Scoring PPL1 DA neurons in 2-day-old (**d**) and 30-day-old (**e**) flies subjected to Brm- or BVR-related genetic manipulations. Genetic manipulations included TH-Gal4 driving overexpression of wide type Brm (Brm wt), a dominant-negative allele of Brm (Brm^DN^), dBVR OE (Bvr II and Bvr III), Bvr II + Brm^DN^ and induction of Brm RNAi, BVR RNAi or Bvr II + Brm RNAi with w- and Luc RNAi flies as the control. The genotypes of experimental flies are provided in the supplementary appendix. Overexpression of dBVR resulted in DA neuronal loss in the aged fly brains when compared with controls (*Bvr* II, 10.37±1.03; *Bvr* III, 10.70±0.67; and control, 11.77±0.42). n > 20 for each data point. **f** Progressive elevation of Brm protein levels (GFP signal) in the brains of four PD model flies in comparison to the control. Representative whole-mount fluorescence images of fly brains are provided, with quantifications of the Brm protein level in young (3-day-old), middle-age (15-day-old) and aged (30-day-old) flies shown below (n > 5). *indicates Mann-whitney *P* < 0.01. Scale bar, 10 μm.

*Drosophila* genetic manipulations were then carried out to dissect the roles of Brm and dBVR. When Brm RNAi-mediated down-regulation [22] or ectopic supply of a dominant negative allele of Brm (Brm^DN^) was induced specifically in DA neurons, no change in the number of PPL1 DA neurons was observed, neither was the overexpression of wild type Brm (Fig. 2d-e). However, pan-neuronal (*elav*-Gal4 driver) overexpression or down-regulation of Brm led to early lethality. Meanwhile, we generated two UAS-dBVR transgenic fly strains, Bvr II and Bvr III, which enabled overexpression of dBVR (Additional file 1: Figure S2). Specific overexpression of dBVR in DA neurons resulted in DA neuronal loss in the aged fly brains as compared with controls, while an available dBVR RNAi line did not show any effects (Fig. 2d-e, Additional file 1: Figure S2). Remarkably, ectopic supply of Brm^DN^ suppressed the progressive DA loss caused by BVR overexpression (Fig. 2d-e). The rescuing effect was not owing to titration of UAS-mediated overexpression (Additional file 1: Figure S3).

### Genetic manipulations of Brm or BVR modulated DA degeneration in common *Drosophila* PD models

We then examined how Brm and dBVR genetically interact with known PD genes. Four previously reported *Drosophila* PD models were successfully reproduced in our laboratory [23, 24]. Given that the homozygous Parkin null alleles were found to be unhealthy in our laboratory, we used Parkin RNA interference (RNAi) flies instead. These four *Drosophila* PD models were abbreviated as αSyn (A30P), Lrrk2 (I1915T), Parkin, Pink1 (Pink1 mut or Pink1 RNAi) PD models respectively (see Methods & fly genotypes listed in the Additional file 1: appendix). Consistent with previous findings, age-dependent progressive degeneration was mild but statistically significant, as evidenced by the decreased number of PPL1 DA neurons in 30-day-old flies compared with age-matched controls. In contrast, young 2-day-old PD flies displayed no DA neuronal loss (Additional file 1: Figure S4).

We found that Brm∷GFP level exhibited an age-dependent progressive elevation in the PD fly brains compared with controls (Fig. 2f). We next addressed how Brm is up-regulated in the degenerative dopaminergic neurons. One possibility is there could be a link between Brm activity and oxidative stress, which has been widely believed to be a common pathogenic factor in PD. To this end, we monitored the oxidative stress indicated by the ROS fluorescent dye (DCF-DA) (Additional file 1: Figure S5) or the reduction-oxidation-sensitive GFP (roGFP) (Additional file 1: Figure S6-S7). We also evaluated the level of anti-oxidant response in the brains of PD model flies using GstD-GFP as a reporter [25] (Additional file 1: Figure S8). To our surprise, unlike Brm which was pronouncedly elevated at 15d AE during the disease progression in all four PD models, no substantial increase of brain oxidative stress reporting signal was detected until 15- 20d AE in the brains of PD flies except for the αSyn detected by roGFP (Additional file 1: Figure S6-S7). Therefore, it implicates that Brm up-regulation in PD flies might not be largely due to the change of DA neuronal oxidative stress.

Remarkably, when Brm RNAi was introduced into the four different PD model flies, significant suppression of PPL1 DA neuronal loss was detected in the aged flies (Fig. 3a-d). As a control, an introduction of irrelevant Luc RNAi did not exhibit such inhibitory effects (Fig. 3a-d). Accordingly, overexpression of Brm^DN^ fully prevented the progressive PPL1 DA neuron degeneration in all four PD fly models, although overexpression of Brm in the PD context did not induce further neuronal loss (Fig. 3a-d, Fig. 2e). The observed rescuing effects did not occur during the developmental stage, since the inhibition of Brm alone did not increase the number of DA neurons more than normal in the young flies (Fig. 2d, Additional file 1: Figure S9a). Neither was the rescuing effect due to titration of UAS-mediated overexpression (Additional file 1: Figure S3).

**Fig. 3.**
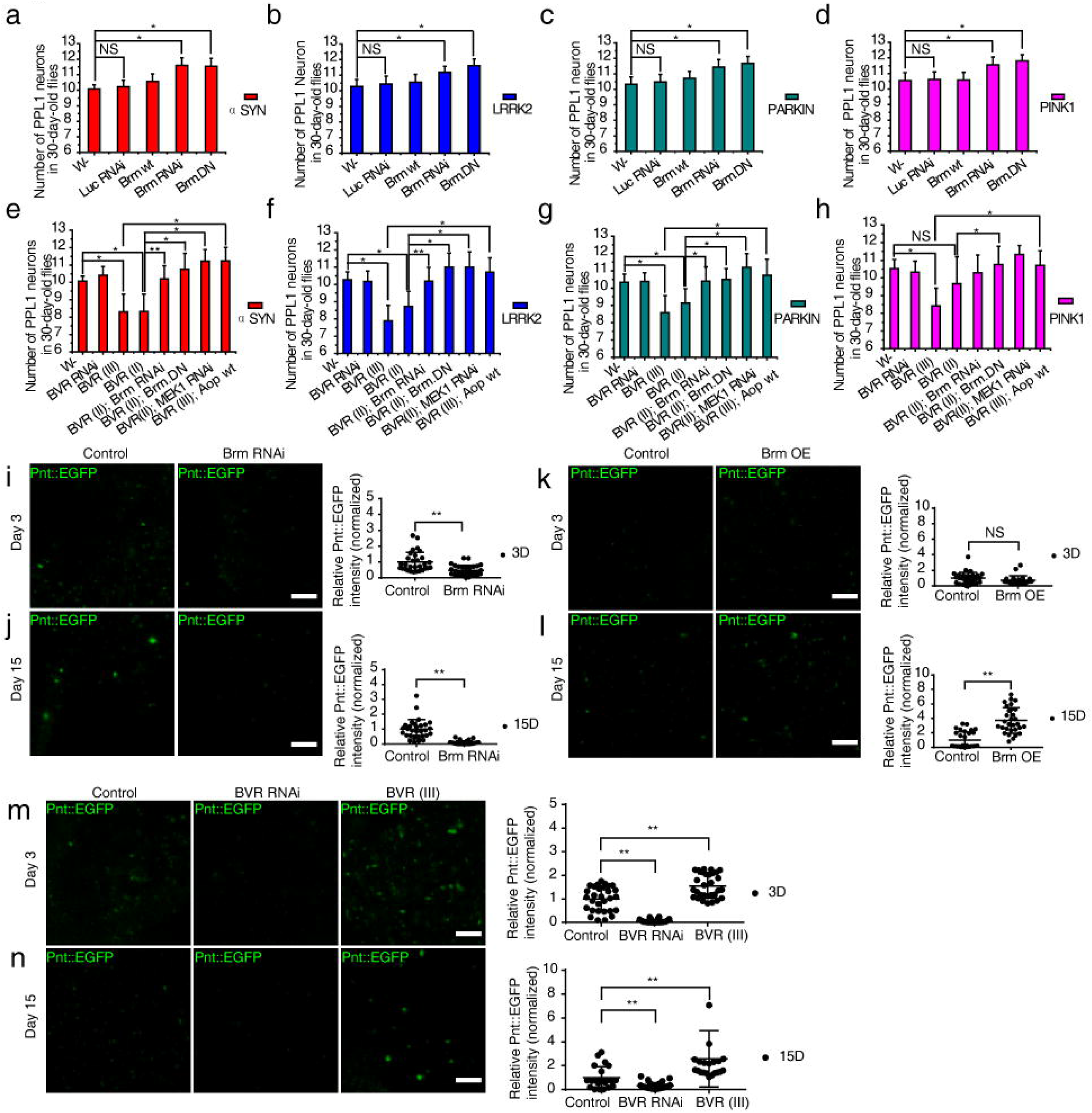
Intervening Brahma or BVR modulated dopaminergic (DA) degeneration in multiple established *Drosophila* PD models. **a–d** RNAi knockdown of Brm prevented DA neuronal loss in four different forms of parkinsonian DA degeneration: αSyn (**a**), Lrrk2 (**b**), Parkin RNAi (**c**), and Pink1 mut (**d**) models. DA degeneration was revealed in the neuronal loss in the PPL1 DA neuron clusters (n > 20). Suppression of Brm by overexpression of the dominant-negative of Brm (Brm^DN^) rescued DA neuronal loss in all four PD fly models (n > 20). **e–h** Overexpression of dBVR aggravated the progressive DA neuronal loss in aged four PD model flies (αSyn, 8.31±1.01; Lrrk2, 7.89±0.90; Pink1 mut, 8.42±1.00; Parkin RNAi, 8.58±1.00), which could be suppressed by MEK RNAi or Aop^wt^ overexpression in all four PD models, the same effects were also found with Brm^DN^ over-expression (n > 20). **i–n** Brm and dBVR positively correlated with the ERK-ETS signaling level indicated by the fusion protein Pnt∷EGFP, which is a transgene to monitor the endogenous protein expression level of Pnt, a reporter for MEK-ERK signaling activity. Representative whole-mount fluorescence images of brains of different ages are shown, with the corresponding quantification of Pnt protein level displayed on the right-side panels (n > 5). The genotypes of experimental flies are provided in the supplementary appendix. *indicates Mann-whitney *P* < 0.05, **indicates *P* < 0.01. NS, not significant. Scale bar, 10 μm.

On the other hand, overexpression of dBVR aggravated the progressive DA neuronal loss in aged four PD model flies, while dBVR RNAi did not mitigate the degeneration (Fig. 3e-h, Additional file 1: Figure S9b). Importantly, the aggravation caused by dBVR overexpression could be fully rescued through the addition of Brm^DN^ in all four PD model flies, and in three PD models (except the *PINK1* deficiency model) through Brm RNAi (Fig. 3e-h). The rescuing effect was not owing to titration of UAS-mediated overexpression (Additional file 1: Figure S3). Collectively, these results demonstrated that inactivation of Brm protects DA neurons from age-dependent degeneration in a variety of pathogenic genetic backgrounds.

### Prolonged over-activated MEK-ERK-ETS signaling in multiple PD fly models

We then investigated the possible mechanisms through which Brm and dBVR could affect DA degeneration. An Ingenuity Pathway Analysis (IPA) revealed the connection of SMARCA4/Brm with ERK signaling pathway (Fig. 1b), as was supported by a previous study [22]. On the other hand, hBVR has been previously suggested as an ERK activator in HEK293A cells [26]. Activator *Pointless* (Pnt) and repressor *Anterior open* (Aop) are two downstream antagonizing players of the MEK-ERK signaling and both belong to E-twenty six transcription factors [27]. We used the reporter fly line of Pnt∷EGFP [27] to monitor the MEK-ERK activity in the fly brain and found that both Brm RNAi and dBVR RNAi resulted in reduced MEK-ERK activity in fly DA neurons, while overexpressing either Brm or dBVR up-regulated the signaling (Fig. 3i-n). Conversely, DA-neuron-specific knock-down of *Drosophila* MEK (*Dsor1*) by RNAi [28] or overexpression of a wild type allele of the negative regulator of ERK pathway, Aop^*[wt]*^, suppressed the aggravation of progressive DA degeneration induced by dBVR overexpression in four PD fly models (Fig. 3e-h); Aop^*[wt]*^ overexpression also prevented DA degeneration caused by dBVR overexpression alone (Additional file 1: Figure S10). These rescuing effects were not due to titration of UAS-mediated overexpression (Additional file 1: Figure S3).

We then examined whether over-activation of ERK-ETS was prevalent in those common PD fly models. Remarkably, we observed increased phosphorylated ERK (pERK) levels in all four PD model fly brains (Fig. 4a-b), which were concordant with sustained up-regulation of Pnt (Fig. 4c-e).

**Fig. 4.**
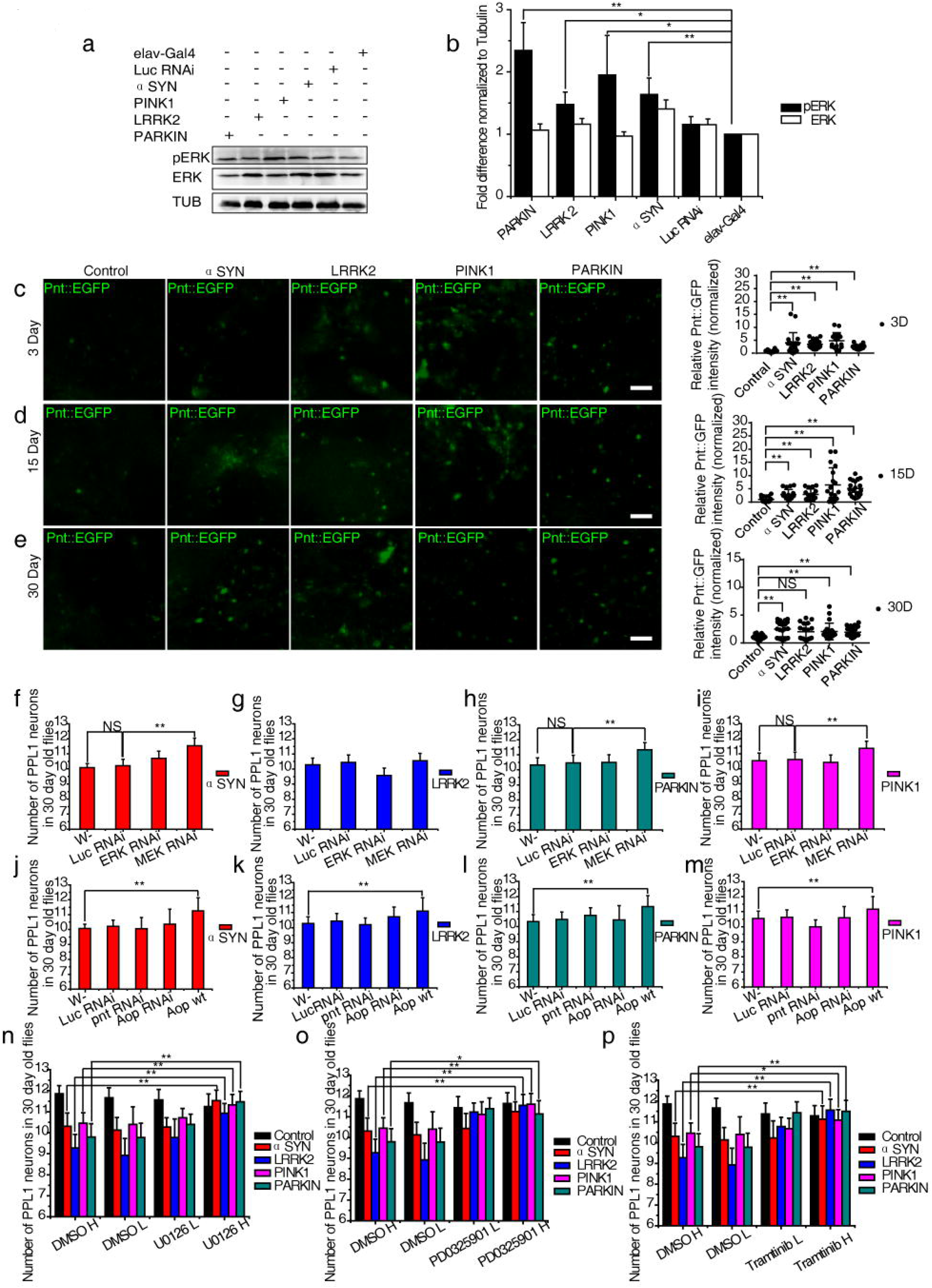
Attenuation of MEK-ERK-ETS signaling activity prevented dopaminergic (DA) degeneration in *Drosophila*. **a, b** The ERK phosphorylation level was significantly enhanced in all four PD model fly brains with *elav*-Gal4 as the driving line. Luc RNAi was used as a control for PINK1RNAi and Parkin RNAi experiments. *indicates Student *t*-test *P* < 0.05, **indicates *P* < 0.01. **c–e** Sustained elevated Pnt level in PD fly brains indicated prolonged over-reactive MEK/ERK signaling. Pnt∷EGFP was used and quantified as in Figure 3. Representative whole-mount fluorescence images of brains of different ages are shown. **f–i** Knockdown of MEK prevented the deterioration of the PPL1 DA neurons in three PD fly models: αSyn, 11.55±0.5 (**f**); Parkin RNAi, 11.33±0.47 (**h**); Pink1 mut, 11.36±0.48 (**i**); and control, 11.67±0.22. **j–m** Overexpression of Aop^wt^ suppressed the degeneration of the PPL1 DA neurons in all four PD fly models: αSyn, 11.25±0.89 (**j**); Lrrk2, 11.1±0.88 (**k**); Parkin RNAi, 11.33±0.72 (**l**); Pink1 mut, 11.17±0.83 (**m**); and control, 11.53±0.32. **n–p** Oral administration of three MEK inhibitors suppressed DA degeneration in *Drosophila* respectively. Two concentrations of either U0126 (**n**) (low [L]: 1 μg/mL; high [H]: 10 μg/mL), PD0325901 (**o**) (L: 1 μg/mL; H: 10 μg/mL), or Trametinib (**p**) (L: 1.6 μM; H: 16 μM) were applied (n > 20). The genotypes of experimental flies are provided in the supplementary appendix. *indicates Mann-whitney *P* < 0.05, **indicates *P* < 0.01. NS, not significant. Scale bar, 10 μm.

### Attenuation of MEK-ERK-ETS activation prevented DA degeneration in multiple *Drosophila* PD models

We next examined whether directly reducing MEK-ERK-ETS activation was sufficient to prevent those common forms of DA degeneration. MEK RNAi fully rescued PPL1 DA neuronal loss in three 30-day-old PD model flies (Fig. 4f-I, Additional file 1: Figure S11a-b) and the rescue was not owing to titration of UAS-mediated overexpression (Additional file 1: Figure S3). In contrast, *Drosophila* ERK (*rolled* or *rl*) RNAi [28] exerted no apparent rescuing effects, suggesting that it was the activated fraction of ERK (pERK), not the abundance of the ERK protein, that caused the neurotoxicity. In parallel, when MEK RNAi or ERK RNAi was induced alone specifically in DA neurons, no changes in the PPL1 DA neurons were observed when compared with age-matched controls, suggesting that the rescuing effects of MEK RNAi was epistatic, but not due to simple addition (Additional file 1: Figure S11a). Overexpression of constitutively-active ERK led to early larval lethality before eclosion. We further found that the DA-neuron-specific overexpression of a constitutively active form of Pnt (*Pnt*^*[P1]*^) led to early lethality before eclosion too. Nevertheless, DA-neuron-specific RNAi knock-down of Pnt [28] did not lead to a rescue or mitigation, probably because the manipulation was not potent enough to significantly disrupt the ERK activity. Remarkably, DA-neuron-specific overexpression of the negative regulator, *Aop*^*[wt]*^, completely blocked the aging-related PPL1 DA neuronal loss in all four PD model flies (Fig. 4j-m, Additional file 1: Figure S12a, c), and the rescue was not owing to titration of UAS-mediated overexpression (Additional file 1: Figure S3). No further aggravation was observed upon DA-specific RNAi [28] knock-down of *Aop* (Fig. 4j-m, Additional file 1: Figure S11a, S11c). Similar to Pnt, DA-neuron-specific overexpression of a constitutively active form of AOP (*AOP*^*[CA]*^) led to early lethality before eclosion, and overexpression of *Aop*^*[wt]*^ or Aop RNAi alone resulted in mild DA neuronal loss (Additional file 1: Figure S11a). Based on all the above evidences, we conclude that there might be a delicate range of the ERK-ETS signaling strength that is beneficial for the maintenance of DA neurons, while deviation from that range such as prolonged over-activation could be rather detrimental.

### Pharmacological inhibition of ERK signaling suppresses DA neuronal loss in multiple *Drosophila* PD models

To examine the MEK/ERK signaling pathway as a drug target for intervening in DA degeneration, we started with the MEK1 inhibitor, U0126. The effective inhibitory dose was first determined in flies subjected to 7 days of drug feeding (Additional file 1: Figure S12a-b). After continuous oral supplementation of U0126 (10 μg/ml) in adult flies for 30 days, DA neuronal loss in PPL1 clusters were blocked effectively in all four PD fly models (Fig. 4n). Exposure to another MEK1 inhibitor, PD0325901 (10 μg/ml), also completely blocked PPL1 DA neuronal loss (Fig. 4o). We further tested Trametinib, another potent and highly specific MEK1 inhibitor, an FDA-approved drug for the treatment of melanoma [29]. With the optimal feeding concentration of 16 μM of Trametinib (Additional file 1: Figure S12c-d) [28], PPL1 DA neurons were fully protected from degeneration (Fig. 4p). No global brain or motor behavioral abnormalities were detected with all these drug treatments. Taken together, our data demonstrated that the MEK-ERK pathway could be a valid drug target to revert DA degeneration and illustrated that oral administration could be a promising pharmacological intervention.

### Drug inhibition of ERK activity ameliorated mitochondrial defects in *pink1*^−/−^ HeLa cells

In humans, *pink1*-deficiency leads to mitochondrial defects [30]. To determine whether our findings were relevant to human pathology, we generated a *pink1*^−/−^ HeLa cell line, using the CRISPR/Cas9 technology (Fig. 5a) [31]. Enhanced phosphorylation of ERK and MEK1 was observed in the *pink1*^−/−^ HeLa cells, indicating an aberrant MAPK signaling over-activation (Fig. 5b). Treatment with the MEK1 inhibitor, PD0325901, effectively mitigated multiple mitochondrial defects in the *pink1*^−/−^ cells, such as reduced mitochondrial membrane potential, lowered mitochondrial contents, and abnormal network interconnectivity (Fig. 5c-d). In agreement with our results, MEK1 inhibitor has been shown to reverse PD-associated phenotypes induced by pathological *LRRK2* alleles in cultured human iPS derived neurons [32].

**Fig. 5.**
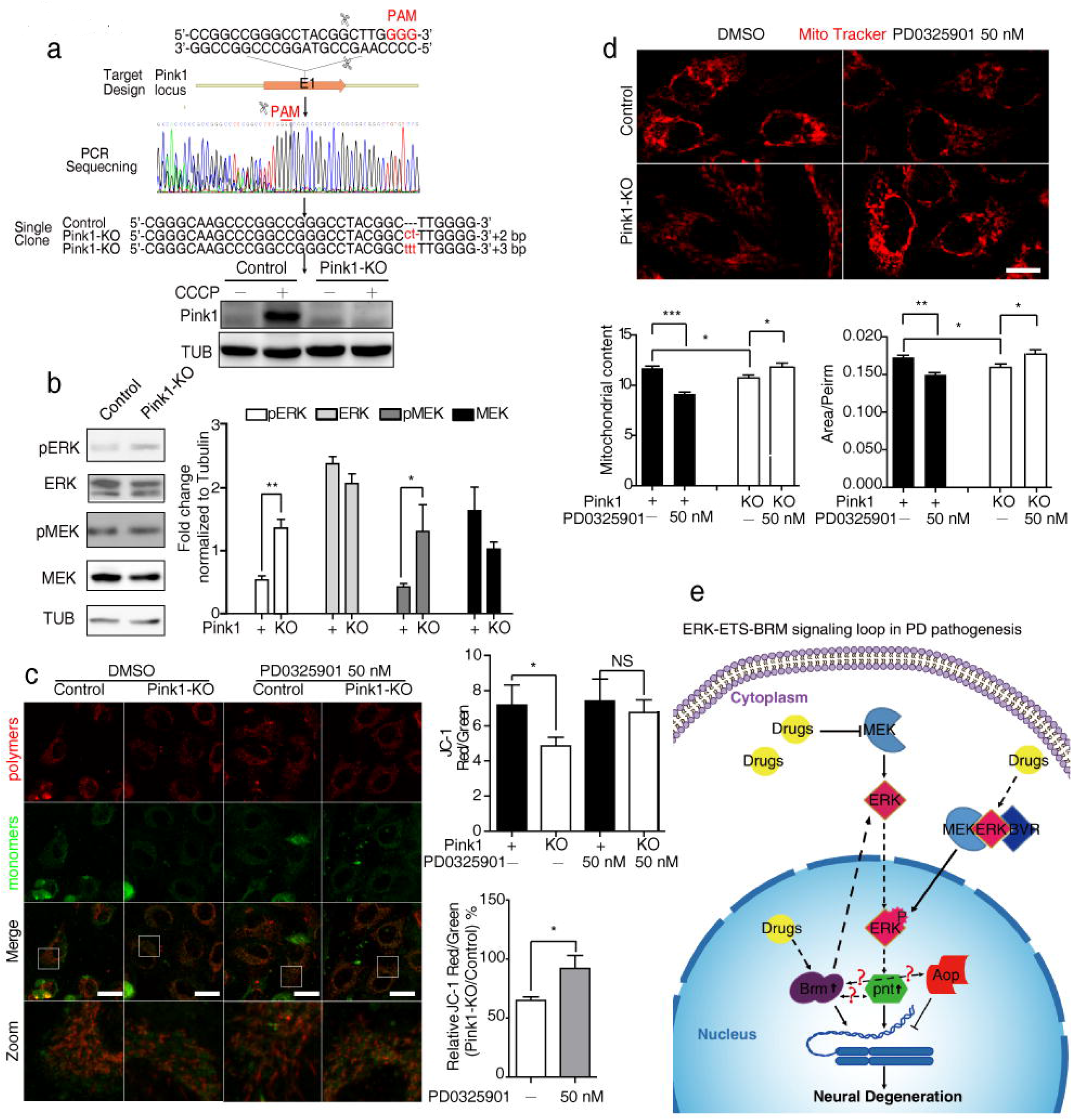
Drug inhibition of MEK-ERK signaling ameliorated mitochondrial defects in *pink1^−/−^*HeLa Cells. **a** CRISPR/Cas9 to establish a *pink1*^−/−^ HeLa cell line. Schematic overview of the strategy used to generate the *pink1* knock-out allele, Sanger sequencing was used to identify the targeted mutations, and western blot analysis verified the loss of PINK1 protein. Control or *pink1* KO HeLa cells, untreated (−) or treated (+) with the protonophore m-chlorophenylhydrazone (CCCP) were used; CCCP was used to prevent PINK1 protein from degradation [41]. **b** Over-activated MEK/ERK signaling in *pink1*^−/−^ HeLa cells. **c** Inhibition of MEK-ERK signaling by PD0325901 rescued mitochondrial membrane potential (MMP) defects in *pink1^−/−^* HeLa cells. The mean ratio of the JC-1 dye intensity in red channel to green channel is shown as mean ±standard deviation (s.d.) to quantify MMP. Relative MMP between *pink1*^−/−^ and control is also shown. **d** Inhibition of MEK/ERK signaling improved mitochondrial content and mitochondrial interconnectivity in *pink1*^−/−^ HeLa cells. Mitochondria were stained with MitoTracker-Red. The mean mitochondrial content and mitochondrial interconnectivity index (Area/Perim) are shown as mean±s.d.. The drug solvent used was DMSO, and equivalent amounts of DMSO were used in parallel to the drug as a treatment control. Representative confocal images and quantifications are shown. *indicates Mann-whitney *P* < 0.05, **indicates *P* < 0.01, ***indicates *P* < 0.001. NS, not significant. Scale bars, 10 μm. **e** A mechanistic model of the signaling loop in PD pathogenesis is proposed and the potential drug intervening points are illustrated.

## DISCUSSION

In summary, we identified a subset of 32 novel PD-associated genes, which were highly enriched in aging and neural disorders. The roles of two candidate genes SMARCA4/Brm and BLVRA/BVR were validated *in vivo*. The activity of ERK-ETS signaling, as a common effector for SMARCA4/Brm and BVR, was found to be also elevated in different genetic forms of *Drosophila* PD models. Thus, we have discovered a potential convergent PD pathogenesis pathway. Our finding also underscores the important role of epigenetic regulators and reveals a novel epigenetic target besides HDACs and DNA methyltransferases (DNMTs) for the therapeutic interventions of aging-related disorders, including PD. Remarkably, the *in vivo* genetic manipulations in our studies were specifically restricted in the dopaminergic neurons thanks to the GAL4-UAS binary expression system in *Drosophila*, highlighting the cell autonomous impacts of the target genes. In fact, genetic modifications of vulnerable dopaminergic neurons *per se* may be valuable for cell/genetic therapy in PD patients in the future.

Brm was found here to be progressively up-regulated in the aging brains of PD fly models. One possibility is through a genetic imprinting response to elevated calcium level triggered by aberrant neuronal activities or calcium metabolism [33]. Alternatively, Brm could be activated through NF-kB mediated inflammatory responses that are well recognized in the development of neurodegenerative diseases [34]. Besides, there is a clue that Brm functions downstream of of Hippo pathway and plays an important role in a feedback loop between Crumb and Yorkie in this pathway [35]. On the other hand, in our unpublished data, we did observe the differential expression of other oxidoreductases (e.g., P450, sulfiredoxin, phenoloxidase) at the early-middle stage of PD. Therefore, it remains possible that Brm might directly interact with Keap1/Nrf to elicit a program of sequential cellular responses to oxidative stress in DA neurons.

Inactivation of Brm was shown here to prevent dopaminergic neurons from degeneration. One putative route was through modulating the MAPK/ERK signally activity. Accordingly, Brm was shown to directly interact with Dsor1 (MEK1) and promotes EGFR-Ras-MAPK signaling activity [22, 36]. Nevertheless, the possibility could not be ruled out that direct interactions exist between Brm with transcriptional factors downstream of MAPK/ERK signaling, such as Pnt or Aop, constituting a positive auto-regulation feedback loop. Alternative mechanisms await further investigation to elucidate the effects of Brm.

Our finding that Brm inactivation protects DA degeneration seems to be at odds with the positive roles of Brm in the neural development [15, 16, 33]. However, previous studies have shown SWI/SNF complexes in different tissues at different stages have distinct functions by eliciting context-specific transcription programs. It is very likely that Brm may need to collaborate with other SWI/SNF components and DA neuronal specific transcription factors to regulate the expression of target genes in DA neurons. Such possibility can help to explain why we did not observe substantial pathogenic effect by overexpressing Brm in the *Drosophila* DA neurons. Dosage and non-linear effect might also contribute to the insufficiency. Nevertheless, future works should clarify the complex role of Brm in the aging neurons.

Different mitogen-activated protein kinase (MAPK) pathways have been linked to aging and aging-related disorders, e.g., different forms of PD, especially the Jun amino-terminal kinases (JNK) and p38 MAPK pathways. However, the role of MEK-ERK is surprisingly elusive and somewhat controversial [32, 37, 38], very likely owing to the fact that most previous studies were performed *in vitro* resulting in contradictory results. Our results thus clarify the hitherto controversial role of MEK-ERK activation in age-related PD pathogenesis by concluding that chronic and prolonged over-activation of MEK-ERK beyond a beneficial range results in DA neurotoxicity.

Three compounds that inhibited MEK-ERK signaling also ameliorated DA degeneration in all PD fly models. Interestingly, Trametinib has been shown to extend lifespan [28], while targeted inhibition of ERK signaling prevents spinocerebellar ataxia type 1 in *Drosophila* and mice models [39]. To our knowledge, this manuscript represents the first report of compounds that are effective in preventing DA degeneration *in vivo* in the four most common genetic forms of PD, compared to a previous report of Rapamycin [40]. MEK-ERK-ETS and mTOR/4E-BP pathways might converge on common downstream effectors such as mitochondrial activity/quality control, proteostasis, autophagy, oxidative stress response, DNA damage repair, and DNA-chromatin modifications, all of which are important aspects during aging. It is conceivable that cocktail strategies using both lines of inhibitors might eventually balance symptoms mitigation and side effects.

## CONCLUSIONS

By combining bioinformatics analysis of large-scale human transcriptomic data and a small-scale genetic screening using *Drosophila* disease models, we identified two novel PD-associated genes SMARCA4 and BLVRA and disclosed a potential common pathogenic PD pathway, i.e. SMARCA4-ERK-ETS signaling axis. We unprecedentedly demonstrated that inhibiting the MEK-ERK signaling by three compounds ameliorated progressive dopaminergic degeneration *in vivo* in 4 most common genetic PD models. Due to the high degree of evolutionary conservation in the SWI/SNF, BLVR/HO, and MEK-ERK-ETS pathways, our study may reveal multiple therapeutic entry points to reverse dopaminergic degeneration and extend the life-span in humans (Fig. 5e).

## LIST OF ABBREVIATIONS

PD: Parkinson’s disease
*SMARCA*4: SWI/SNF related, matrix associated, actin dependent regulator of chromatin, subfamily A, member 4
*BLVRA*: biliverdin reductase A

## Acknowledgments

We thank Mu-ming Poo, Bingwei Lu, and Yu Cai for critical comments on the manuscript. We also would like to thank Bingwei Lu, Nancy Bonini, Helena E. Richardson, and Jongkyeong Chung for sharing key fly strains, Wei Wu at Core Facility of *Drosophila* Resource and Technology, SIBCB, CAS for providing fly service and fly stocks, and Tsinghua *Drosophila* Resource Center for providing fly stocks.

## DECLARATIONS

### Funding

This work was supported in part by the National Natural Science Foundation of China (No. 31071296 & 31771410), National Key Basic Research Program of China (973 project, No. 2015CB352006), Province Min-River Scholar Grant (No. XRC-0947), the National Natural Science Foundation of China (No. 31070877), Startup Funds from Fuzhou University (Grant #XRC-1465), the National Natural Science Foundation of China (No. 31601894), Fujian Natural Science Foundation (Grant #2016J01158 & 2017J0106), Education and Scientific Research Projects for Young Teachers in Fujian Province (JAT160071), and United Arab Emirates University Research Grant (31M090).

### Availability of data and material

The gene expression datasets in this study are available from NCBI Gene Expression Omnibus (GEO) at https://www.ncbi.nlm.nih.gov/geo/.

### Author Contributions

Y.Y. conceived the project. Y.Y. and K.H. designed and supervised the experiments. J.Z. and K.H. performed gene co-expression analysis. Y.Y., L.S., Y.C., W.C., Y.Z., Y.Y., M.X., F.M., Y.R., F.C., M.E.H, S.N., Z.Y., Z.S. and X.Z. performed NCBI GEO dataset search, fly genetics, immunohistochemistry, and imaging experiments. L.S. and X.Z. performed p-element constructions and microinjections. L.S, Y.Z., Y.C., X.Z. and H.Z. performed western blotting and drug treatment experiments. W.C., Z.Z., M.Y., Y.Y., M.Y., and K.C. performed human cells studies. Y.Y., L.S., J.Z., and K.H. wrote the manuscript, with contributions from other co-authors

### Ethics approval and consent to participate

Not applicable.

### Consent for publication

All authors have read and approved the final version of the manuscript for publication.

### Competing financial interests

The authors declare no competing financial interests with patent applications having been filed relating to this work.

## ADDITIONAL FILES

### Additional files 1

**Figure S1**. The subcellular localization of Brm∷GFP.

**Figure S2**. Semi qPCR experiments verified the RNAi and overexpression effects of dBVR.

**Figure S3**. Quantitative RT-PCR analysis of dBVR and αSYN to exclude the potential titration problem of UAS-Gal4 system.

**Figure S4**. Establishment of four *Drosophila* PD models.

**Figure S5**. Oxidative stress level indicated DCF-DA in the brains of four PD model flies.

**Figure S6**. Oxidative stress level indicated by tub-mito-roGFP2 in the brains of four PD model flies.

**Figure S7**. Oxidative stress level indicated by UAS-roGFP2 in the PPL1 neurons of four PD model flies.

**Figure S8**. Anti-oxidant response level indicated by GstD-GFP in the brains of four PD model flies.

**Figure S9**. Genetic manipulation of Brahma or BVR modulated dopaminergic degeneration in *Drosophila*.

**Figure S10**. Aop^*[wt]*^ overexpression prevented DA degeneration caused by dBVR overexpression.

**Figure S11**. Genetic manipulation of MEK-ERK-ETS signaling axis modulated dopaminergic degeneration in *Drosophila*.

**Figure S12**. Efficacy of MEK1 inhibitors in fly brains delivered by oral administration.

**Table S1**. Correlation of candidate genes with anchor genes.

**Table S2**. Pearson correlation coefficients (PCC) value between anchors genes and SMARCA4/BLVRA.

**Table S3**. SMARCA4 PD risk SNPs.

**Table S4**. BLVRA PD risk SNPs.

**Table S5**. Oligos used for the experiments.

## Additional files 2

**APPENDIX:** Fly genotype list

