## Supplementary material for "Attenuation of epigenetic regulator SMARCA4 and ERK-ETS signaling suppresses aging-related dopaminergic degeneration": Figure S1-S12, Table S1-S5

**Supplementary Figures**

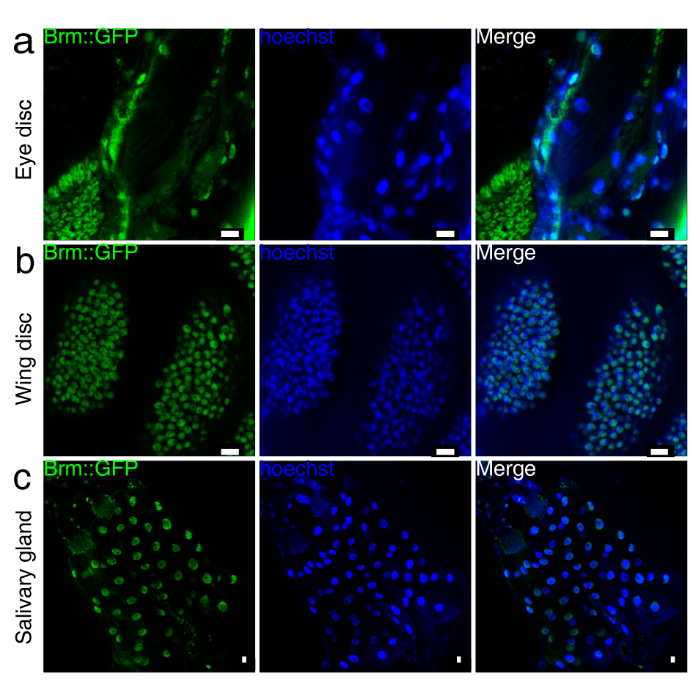

**Figure S1 | The subcellular localization of Brm::GFP.** **a–c,** Representative live images of eye discs (**a**), wing discs (**b**) and salivary glands (**c**) from Brm::GFP larvae. Nuclei were stained with Hoechst in blue. Scale bar, 10 μm.

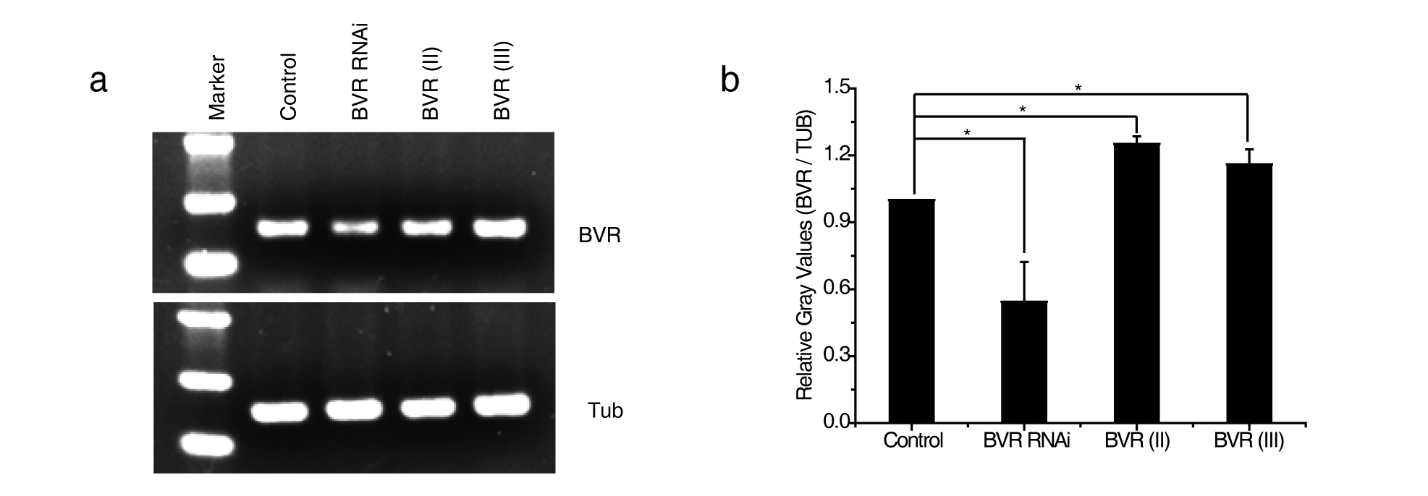

**Figure S2 | Semi qPCR experiments verified the RNAi and overexpression effects of dBVR.** *elav*-Gal4 was used as the driver line. Representative electrophoresis image (**a**) and the quantifications (**b**) were shown. Tubulin served as the internal control. Mean ± SEM was shown (n≥3). * indicates two-tails Student’s t-test with *P* < 0.05.

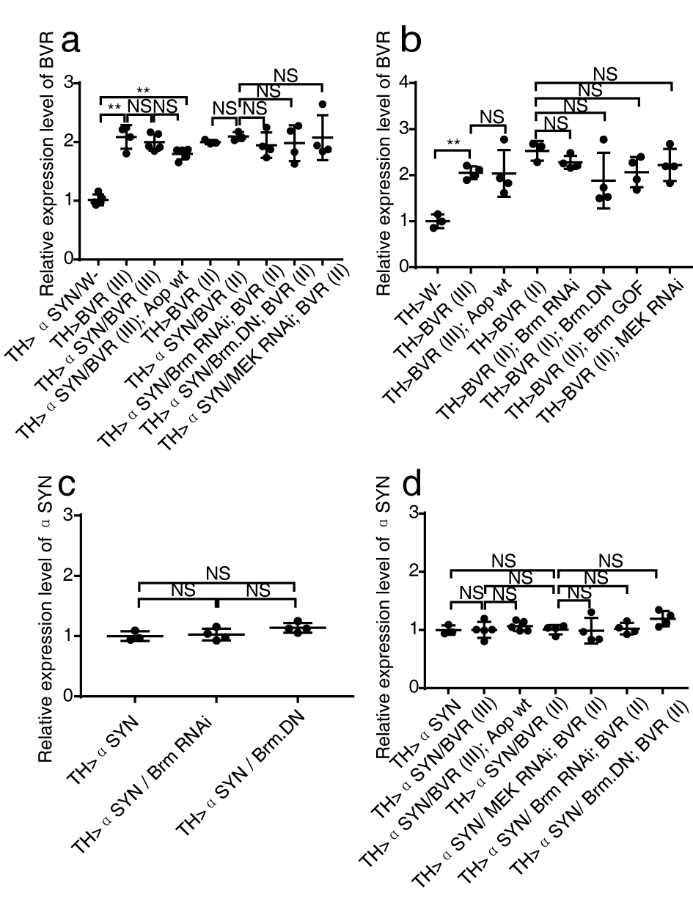

**Figure S3 | Quantitative RT-PCR analysis of dBVR and αSYN to exclude the potential titration problem of UAS-Gal4 system.** **a–b,** Analysis of dBVR expression level in the adult fly brains. Genetic manipulations included TH-Gal4 driving overexpression of αSYN, Bvr III, αSYN +Bvr III, αSYN + Bvr III + Aop^wt^, Bvr II, αSYN + Bvr II, αSYN + Bvr II + Brm RNAi, αSYN + Bvr II + Brm^DN^, αSYN + Bvr II + MEK RNAi (**a**) and TH-Gal4 driving overexpression of Bvr III + Aop^wt^, Bvr II + Brm RNAi, Bvr II + Brm^DN^, Bvr II + Brm^GOF^, Bvr II + MEK RNAi (b). Mean ± SEM (n≥3). **c–d,** Analysis of αSYN expression level in the adult fly brains. Genetic manipulations included TH-Gal4 driving overexpression of αSYN, αSYN + Brm RNAi, αSYN + Brm^DN^ (**c**) and TH-Gal4 driving overexpression of αSYN, αSYN + Bvr III, αSYN + Bvr III + Aop^wt^, αSYN + Bvr II, αSYN + Bvr II + MEK RNAi, αSYN + Bvr II + Brm RNAi, αSYN + Bvr II + Brm^DN^ (**d**). Mean ± SEM was shown (n≥3). ** indicate Mann-whitney with *P* < 0.01. NS, not significant.

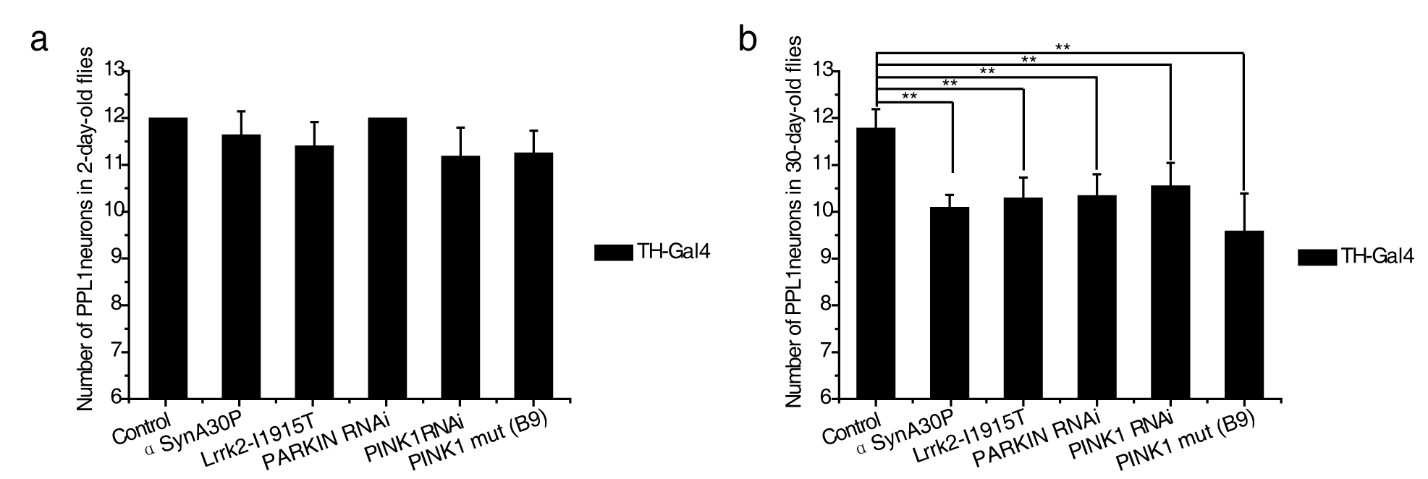

**Figure S4 | Establishment of four *Drosophila* PD models.** **a–b,** Degenerative DA neuronal loss in PPL1 clusters in four *Drosophila* PD models, abbreviated as αSyn, Lrrk2, Parkin, Pink1 (Pink1 null mutant [Pink1 Mut] and Pink1 RNAi) PD models respectively. Control flies were TH-Gal4>w-. PPL1 DA neurons that were marked by anti-tyrosine hydroxylase (TH) antibody were counted. 2-day old (**a**) and 30-day-old (**b**) adult flies were used for analysis, n > 20 for each data point. The genotypes for experimental flies are provided in the supplementary appendix. PPL1 DA neurons were reduced in number in 30-day-old PD flies (αSyn, 10.08±0.28; Lrrk2, 10.28±0.45; Pink1 RNAi, 10.54±0.5; Pink1 mut, 9.57±0.82; and Parkin, 10.33±0.47) compared with age-matched controls (11.77±0.42). ** indicates Mann-whitney with *P* < 0.01.

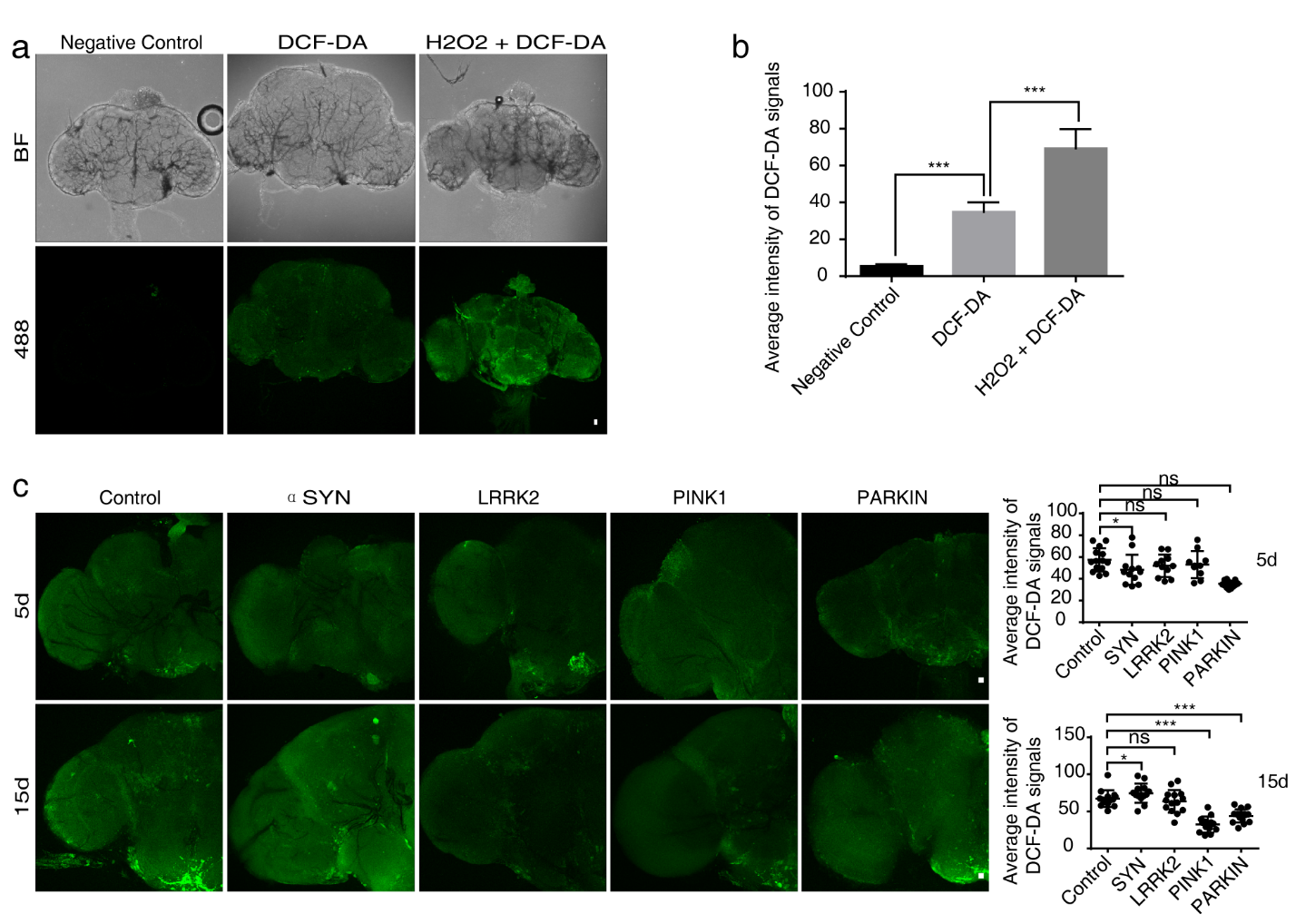

**Figure S5 | Oxidative stress level indicated DCF-DA in the brains of four PD model flies.** **a– b,** Representative brain images of control and 10 μM DCF-DA staining brains with or without 100 μM H2O2 were shown (**a**). Quantification of DCF-DA fluorescent signals (**b**) in *Drosophila* brains were shown. **c,** The DCF-DA staining brains of 5^th^ day AE and 15^th^ day AE in control and four PD model *Drosophila* were analyzed and quantified. Error bars represent the SD. Mann-whitney was performed. ****P*<0.001; ns, not significant. Scale bars, 10 μm.

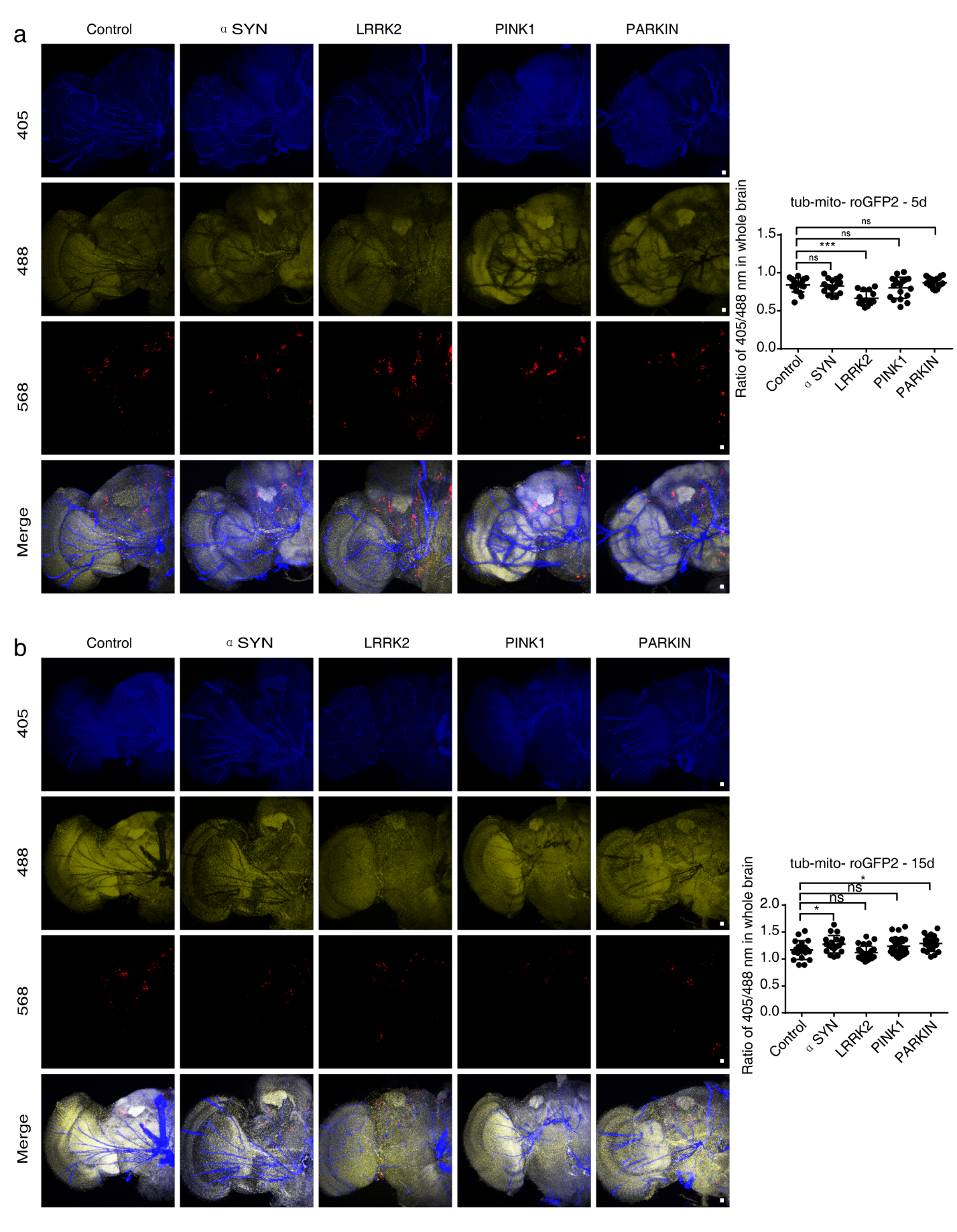

**Figure S6 | Oxidative stress level indicated by tub-mito-roGFP2 in the brains of four PD model flies.** The alphaTub84B regulatory sequences were used to control the pan-expression of mito-roGFP2 in the transgenic reporter flies. Mito-roGFP2 is an oxidant receptor peroxidase-based, mitochondria-localized, fluorescent sensor of hydrogen peroxide oxidation. The oxidative stress was indicated by ratio 405/488 nm of roGFP2 signal. Red fluorescent protein (RFP) was used to label DA neurons. Representative whole-mount fluorescence images of control and PD model fly brains of 5^th^ day AE (**a**) and 15^th^ day AE (**b**) were shown. At least 10 samples were quantified in each experimental group. *** indicates Mann-whitney with *P* < 0.001, NS means ‘not significant’. Scale bar, 10 μm.

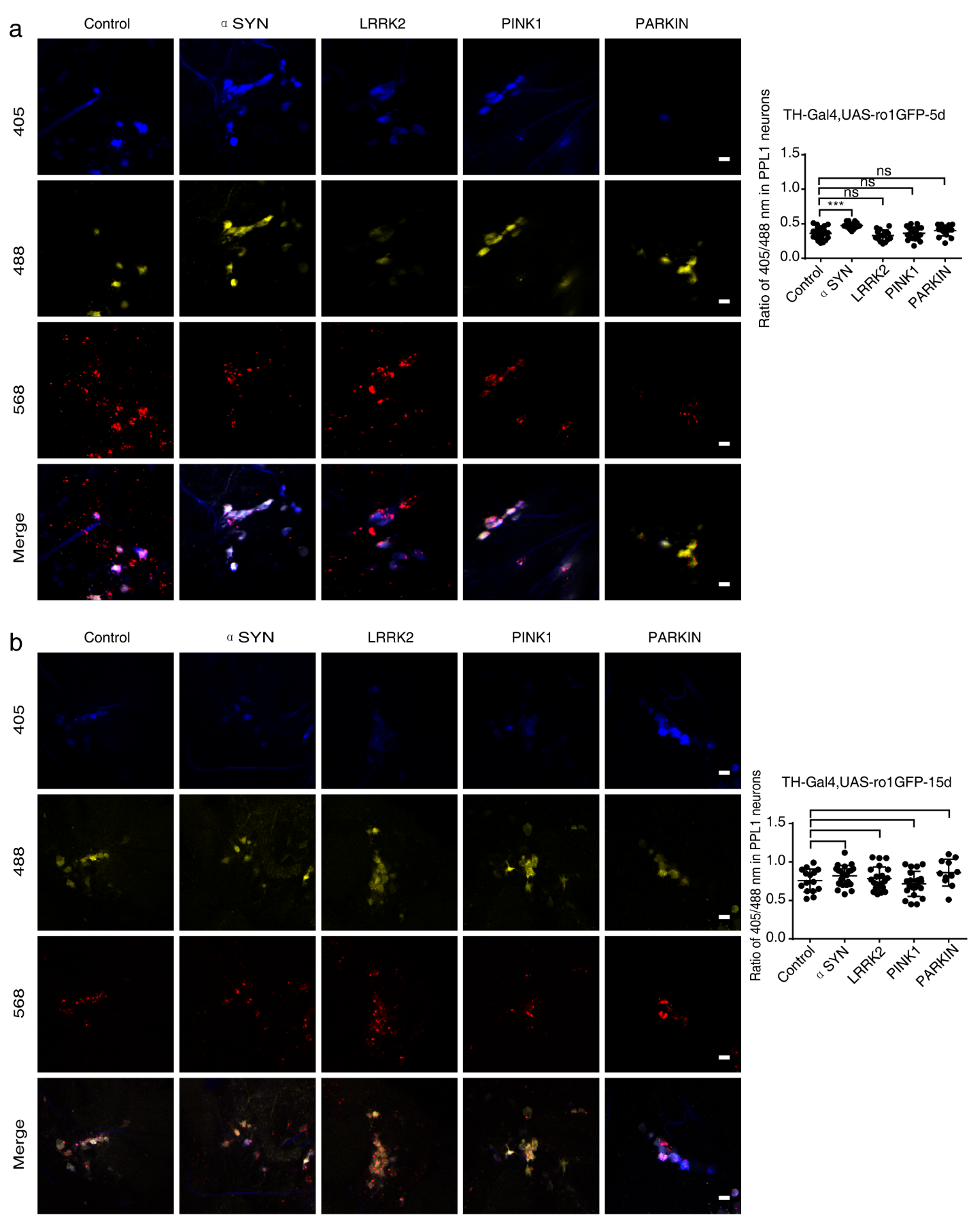

**Figure S7 | Oxidative stress level indicated by UAS-roGFP2 in the PPL1 neurons of four PD model flies.** The fluorescence images show roGFP2 (blue and yellow) fluorescence of PPL1 neurons in fly brains of 5^th^ day AE (**a**) and 15^th^ day AE (**b**). Red fluorescent protein (RFP) was used to label DA neurons. The oxidative stress of PPL1 neurons indicated by ratio 405/488 nm of roGFP2 signal were analyzed and quantified in control and four PD mode flies. At least 10 samples were quantified in each experimental group. *** indicates Mann-whitney with *P* < 0.001, NS means ‘not significant’. Scale bar, 10 μm.

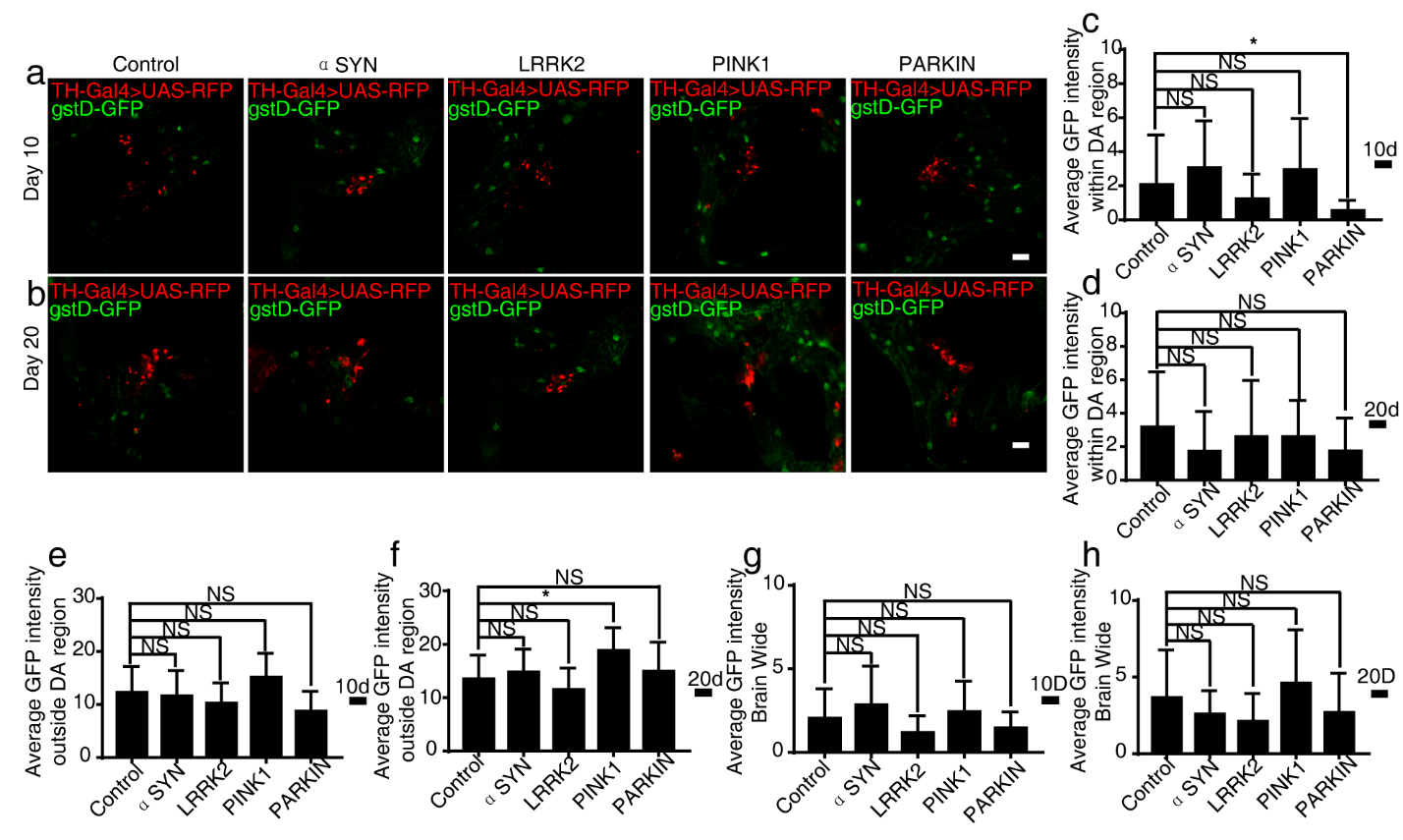

**Figure S8 | Anti-oxidant response level indicated by GstD-GFP in the brains of four PD model flies.** The genomic sequence upstream of the GstD1 gene was used to control the expression of GFP in the transgenic reporter flies [17]. The transcriptional activity of the GstD enhancer indicated by GFP signal can be induced by oxidants and thus can be served an index of oxidative stress. Red fluorescent protein (RFP) was used to label DA neurons. **a–b,** Representative whole-mount fluorescence images of control and PD model fly brains of 10^th^ day AE (**a**) and 20^th^ day AE (**b**) were shown. GstD-GFP signal intensity within the DA region (**c, d**), outside the DA region (**e, f**) and brain-wide (**g, h**) were quantified (n > 5). * indicates two tails Mann-whitney with *P*<0.05, NS means“not significant”. Scale bar, 10 μm.

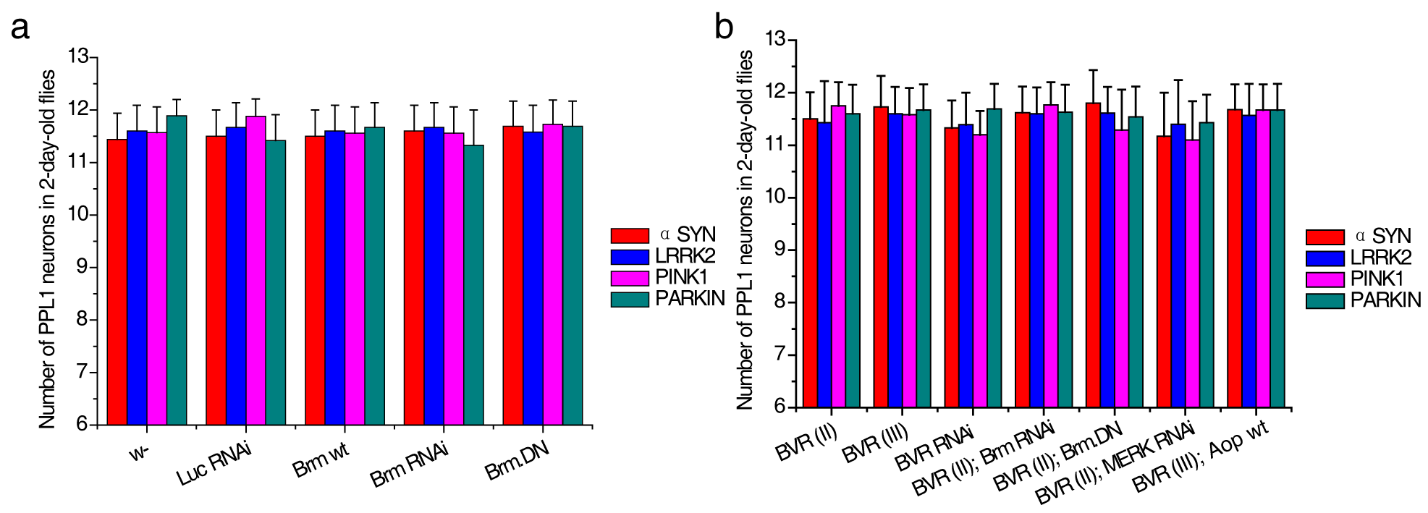

**Figure S9 | Genetic manipulation of Brahma or BVR modulated dopaminergic degeneration in *Drosophila*. a,** Scoring PPL1 DA neurons in 2-day-old flies subjected to Brm-related genetic manipulations. Genetic manipulations included TH-Gal4 driving overexpression of wide-type Brm (Brm wt), a dominant-negative form of Brm (Brm^DN^) and induction of Brm RNAi with w- and Luc RNAi flies as the control. n > 20 for each data point. **b,** Scoring PPL1 DA neurons in 2-day-old flies subjected to dBVR-related genetic manipulations. Genetic manipulations included TH-Gal4 driving overexpression of wide-type BVR (Bvr II or Bvr III), Bvr II + Brm RNAi, Bvr II + Brm^DN^, Bvr II + MEK RNAi, Bvr III + Aop^wt^ and induction of BVR RNAi with w- and Luc RNAi flies as the control. n>20 for each data point.

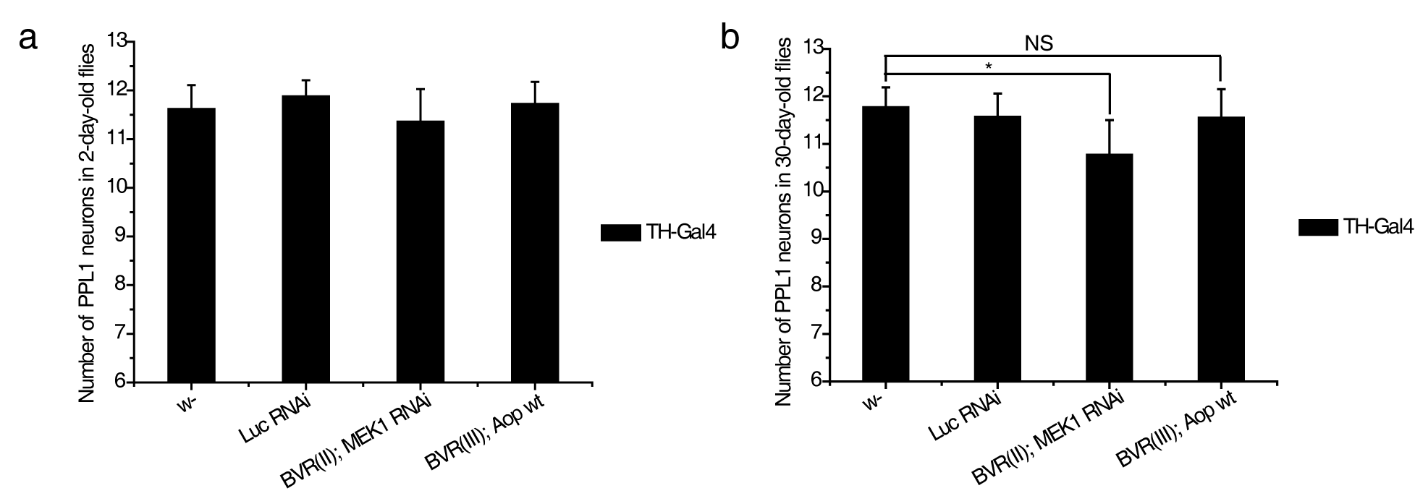

**Figure S10 | Aop*^[wt]^* overexpression prevented DA degeneration caused by dBVR overexpression.** Genetic manipulations included TH-Gal4 driving overexpression of Bvr II + MEK RNAi, Bvr III + Aop^wt^ with w- and Luc RNAi flies as the control. PPL1 DA neurons in 2-day-old (**a**) and 30-day-old (**b**) flies were scored for the phenotype. n > 20 for each data point. * indicates Mann-whitney with *P*<0.05. NS, not significant.

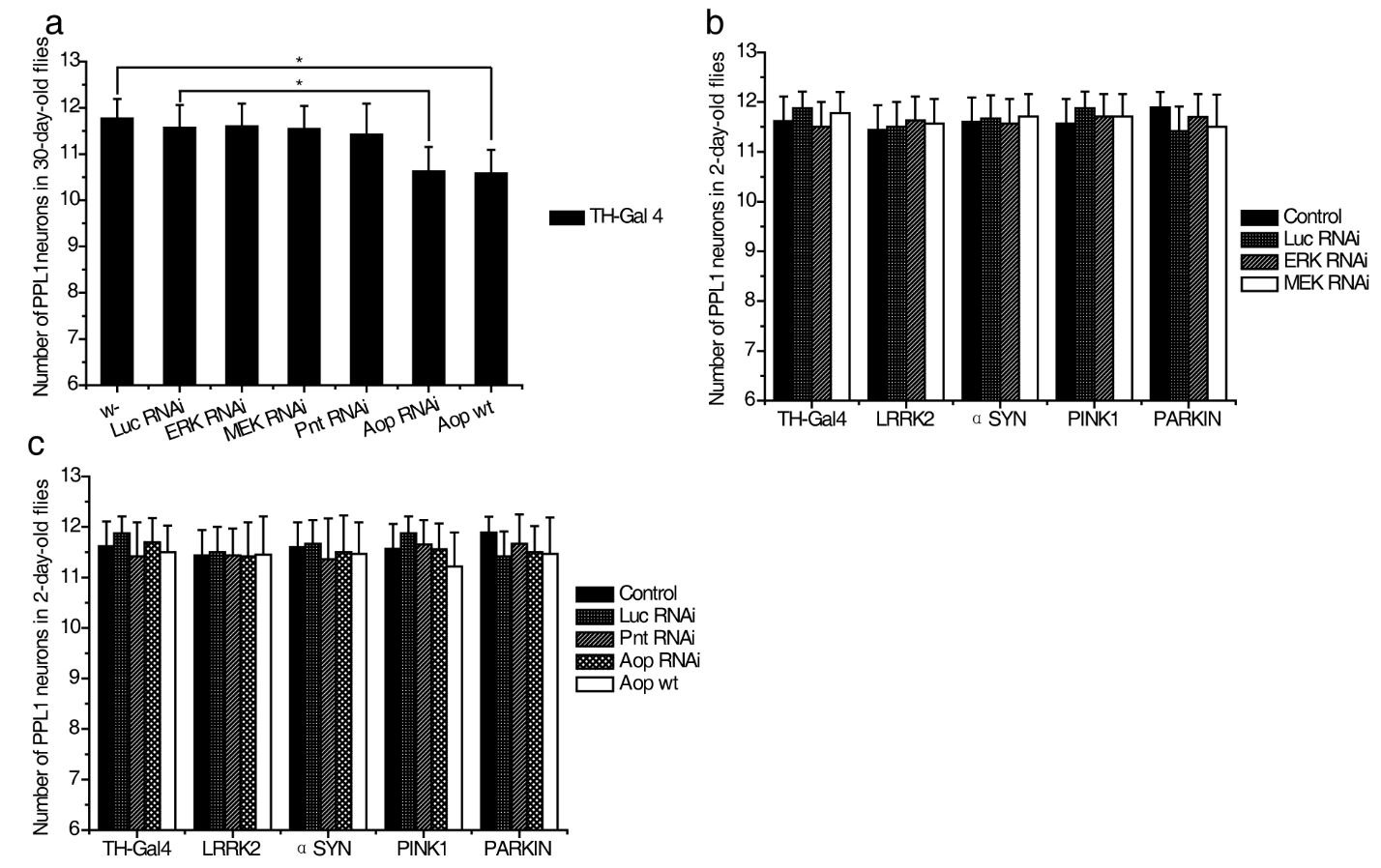

**Figure S11 | Genetic manipulation of MEK-ERK-ETS signaling axis modulated dopaminergic degeneration in *Drosophila*.** **a,** Results of DA neuron-specific genetic manipulations of MEK, ERK, Pnt and Aop. Genetic manipulations included TH-Gal4 driving induction MEK RNAi, ERK RNAi, Pnt RNAi, Aop RNAi and overexpression of Aop, with w- and Luc RNAi flies as the control. PPL1 DA neurons in 30-day-old flies were scored for the phenotype. **b,** Scoring PPL1 DA neurons in 2-day-old flies subjected to MEK or ERK RNAi genetic manipulations with w- and Luc RNAi flies as the control. **c,** Scoring PPL1 DA neurons in 2-day-old flies subjected to Pnt/Aop-related genetic manipulations. Genetic manipulations included TH-Gal4 driving overexpression of wide-type Aop (Aop^wt^), induction of Pnt RNAi and Aop RNAi, with w- and Luc RNAi flies as the control. n > 20 for each data point. *indicates two tails Mann-whitney with *P*<0.05.

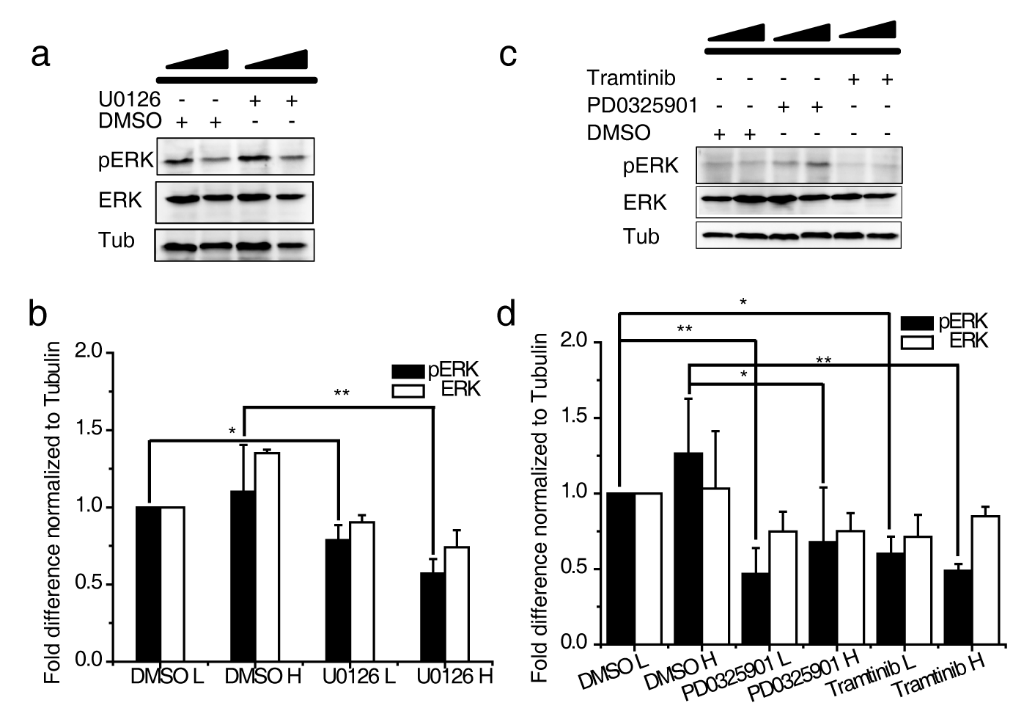

**Figure S12 | Efficacy of MEK1 inhibitors in fly brains delivered by oral administration. a–b,** Inhibitory effect of orally delivered U0126 upon fly brains was validated. Two concentrations of U0126 were applied (L: 1 μg/mL; H: 10 μg/mL). Homogenates of adult fly brains were used for western blot analysis after continuous drug treatment for 7 days. Inactivation of MAPK/ERK was quantified in (**b**) with beta-tubulin as the input control. **c–d,** Effective concentrations of PD0325901 (L: 1 μg/mL; H: 10 μg/mL) or Trametinib (L: 1.6 μM; H: 16 μM) were determined, respectively. DMSO was the solvent and equivalent amount was used in parallel as the drug treatment control. *elav*-Gal4/+ flies were used. Two biological replicates were performed. ** indicates two tails Student’s *t*-test with *P*<0.01, * indicates *P*<0.05.

**Table S1. Correlation of candidate genes with anchor genes**

| **Relativity** | **ATP13A2** | **SCNA** | **HTRA2** | **PARK2** | **LRRK2** | **PARK7** | **PINK1** |
| --- | --- | --- | --- | --- | --- | --- | --- |
| SMARCA4 |  | ※ | ※ |  | ※ | ※ | ※ |
| UBE3A |  | ※ |  |  | ※ | ※ | ※ |
| SNRPN |  | ※ |  |  | ※ | ※ | ※ |
| SLC25A3 |  | ※ | ※ |  |  | ※ | ※ |
| PRDX2 |  | ※ | ※ |  |  | ※ | ※ |
| GNAS | ※ | ※ | ※ |  |  |  | ※ |
| ARF1 | ※ | ※ | ※ |  |  | ※ |  |
| ACTG1 |  | ※ | ※ |  |  | ※ | ※ |
| VCP |  | ※ |  |  |  | ※ | ※ |
| TTC3 |  | ※ |  |  |  | ※ | ※ |
| STMN2 | ※ | ※ |  |  |  |  | ※ |
| RNF187 | ※ | ※ |  |  |  |  | ※ |
| OBSL1 | ※ | ※ |  |  |  |  | ※ |
| OAZ1 |  | ※ |  |  |  | ※ | ※ |
| NSFL1C | ※ | ※ |  |  |  | ※ |  |
| NTRK3 |  |  |  |  | ※ | ※ | ※ |
| MAP2K4 | ※ | ※ |  |  |  |  | ※ |
| LARP1 |  | ※ |  |  |  | ※ | ※ |
| KLC1 | ※ |  |  |  |  | ※ | ※ |
| IGHG1 | ※ |  | ※ |  | ※ |  |  |
| GAPDH |  |  | ※ |  |  | ※ | ※ |
| GAP43 | ※ | ※ |  |  |  |  | ※ |
| EPB41L1 | ※ | ※ |  |  |  |  | ※ |
| DSTN |  | ※ |  |  |  | ※ | ※ |
| DKK3 |  | ※ |  |  |  | ※ | ※ |
| CLTA | ※ |  | ※ |  |  | ※ |  |
| BLVRA | ※ | ※ |  |  |  |  | ※ |
| ATP6V1C1 | ※ | ※ |  |  |  | ※ |  |
| ATP50 |  | ※ | ※ |  |  | ※ |  |
| ARF3 |  | ※ |  |  |  | ※ | ※ |
| AAK1 |  | ※ |  |  | ※ | ※ |  |

Note: ※ represents high correlation between the candidate and the anchor gene.

**Table S2. Pearson correlation coefficients (PCC ) value between anchors genes and SMARCA4/BLVRA**

| HTRA2 | **GDS2519** | **GDS2821** | **GDS3128** | **GSE19587** | **GSE20141** | **GSE20146** | **GSE20153** | **GSE20295** | **GSE20333** | **GDS3129** | **Median** |
| --- | --- | --- | --- | --- | --- | --- | --- | --- | --- | --- | --- |
| SMARCA4 | -0.134385 | 0.401202 | 0.434356 | -0.33275 | 0.778465 | 0.443811 | 0.544739 | 0.940379 | -0.380482 |  | 0.434356 |
| BLVRA | 0.130024 | 0.305576 | 0.514849 | 0.452213 | 0.259953 | 0.350115 | -0.427627 | 0.914044 | -0.267632 |  | 0.305576 |
| LRRK2 |  |  |  |  |  |  |  |  |  |  |  |
| SMARCA4 |  | 0.464852 |  | NA | -0.578301 | -0.4052 | 0.412829 | NA | NA | NA | 0.0038145 |
| BLVRA |  | 0.354546 |  | NA | -0.598422 | 0.406208 | 0.418558 | NA | NA | -0.113976 | 0.354546 |
| PARK2 |  |  |  |  |  |  |  |  |  |  |  |
| SMARCA4 | -0.181832 | 0.448349 | 0.759123 | 0.777099 | 0.217334 | 0.743286 | -0.38967 | 0.952408 | -0.458491 |  | 0.448349 |
| BLVRA | 0.226116 | -0.22987 | 0.73218 | 0.581159 | 0.233454 | -0.531026 | 0.32598 | 0.747416 | -0.077776 |  | 0.233454 |
| PARK7 |  |  |  |  |  |  |  |  |  |  |  |
| SMARCA4 | 0.399054 | 0.653309 |  | 0.696794 | 0.653024 | 0.462393 | 0.387262 | 0.976791 | -0.131253 | NA | 0.5577085 |
| BLVRA | 0.248587 | 0.730552 |  | -0.644057 | 0.811627 | 0.266754 | 0.265855 | 0.960803 | 0.379182 | -0.157136 | 0.266754 |
| PINK1 |  |  |  |  |  |  |  |  |  |  |  |
| SMARCA4 | 0.19246 | 0.608031 | 0.838268 | 0.716564 | 0.9534 | 0.684522 | -0.508087 | 0.932782 | NA |  | 0.700543 |
| BLVRA | -0.292087 | 0.569467 | 0.672501 | -0.574555 | 0.865426 | 0.362951 | -0.489034 | 0.558772 | NA |  | 0.4608615 |
| SNCA |  |  |  |  |  |  |  |  |  |  |  |
| SMARCA4 | -0.285349 | 0.689499 | 0.650606 | 0.698502 | 0.786245 | 0.524943 | 0.426371 | 0.950556 | 0.200241 |  | 0.650606 |
| BLVRA | -0.271254 | 0.525958 | 0.685866 | -0.378715 | 0.724884 | -0.552131 | 0.321484 | 0.924258 | 0.458517 | -0.528703 | 0.458517 |

Note: HTRA2, LRRK2, PARK2, PARK7, PINK1 and SNCA are anchor genes; GDS2519, GDS2821, GDS3128, GSE19587, GSE20141, GSE20146, GSE20153, GSE20295, GSE20333 and GDS3129 are datasets from NCBI Gene Expression Omnibus (GEO). Top 5% PCC are highlighted in yellow.

**Table S3. SMARCA4 PD risk SNPs**

| **[Polymorphism](http://www.pdgene.org/view?gene=SMARCA4)** | **[Location (hg19)](http://www.pdgene.org/view?gene=SMARCA4)** | **Gene** | **[Ethnicity](http://www.pdgene.org/view?gene=SMARCA4)** | **[# Samples](http://www.pdgene.org/view?gene=SMARCA4)** | **[# Studies](http://www.pdgene.org/view?gene=SMARCA4)** | **[Allele contrast](http://www.pdgene.org/view?gene=SMARCA4)** | **1000G CEU** | **1000G CHB+JPT** | **Meta OR (95%CI)** | **I2 (95%CI)** | **[Meta](http://www.pdgene.org/view?gene=SMARCA4)**  [**P-value**](http://www.pdgene.org/view?gene=SMARCA4) |
| --- | --- | --- | --- | --- | --- | --- | --- | --- | --- | --- | --- |
| [rs9105](http://www.pdgene.org/view?poly=rs9105) | [chr19:11169514](http://genome.ucsc.edu/cgi-bin/hgTracks?org=human&hgt.customText=http://www.pdgene.org/tracks?hg=hg19&db=hg19&position=chr19%3A11169264-11169764&hgt.suggest=&pix=800&Submit=submit&hgsid=184845147) | [SMARCA4](http://www.pdgene.org/view?gene=SMARCA4) | C | - | 12 | T vs. C | 0.058 (T) | - | <1 (-) | 0 (-) | <0.05 |
| [rs150323051](http://www.pdgene.org/view?poly=rs150323051) | [chr19:11058880](http://genome.ucsc.edu/cgi-bin/hgTracks?org=human&hgt.customText=http://www.pdgene.org/tracks?hg=hg19&db=hg19&position=chr19%3A11058630-11059130&hgt.suggest=&pix=800&Submit=submit&hgsid=184845147) | [SMARCA4[-12718bp]](http://www.pdgene.org/view?gene=SMARCA4) | C | - | 11 | A vs. G | - | - | <1 (-) | 0 (-) | <0.05 |
| [rs117234045](http://www.pdgene.org/view?poly=rs117234045) | [chr19:11127139](http://genome.ucsc.edu/cgi-bin/hgTracks?org=human&hgt.customText=http://www.pdgene.org/tracks?hg=hg19&db=hg19&position=chr19%3A11126889-11127389&hgt.suggest=&pix=800&Submit=submit&hgsid=184845147) | [SMARCA4](http://www.pdgene.org/view?gene=SMARCA4) | C | - | 11 | A vs. G | 0.008 (A) | - | >1 (-) | 0 (-) | <0.05 |
| [rs118107587](http://www.pdgene.org/view?poly=rs118107587) | [chr19:11134752](http://genome.ucsc.edu/cgi-bin/hgTracks?org=human&hgt.customText=http://www.pdgene.org/tracks?hg=hg19&db=hg19&position=chr19%3A11134502-11135002&hgt.suggest=&pix=800&Submit=submit&hgsid=184845147) | [SMARCA4](http://www.pdgene.org/view?gene=SMARCA4) | C | - | 11 | A vs. G | 0.05 (A) | - | <1 (-) | 0 (-) | <0.05 |
| [rs117819913](http://www.pdgene.org/view?poly=rs117819913) | [chr19:11123091](http://genome.ucsc.edu/cgi-bin/hgTracks?org=human&hgt.customText=http://www.pdgene.org/tracks?hg=hg19&db=hg19&position=chr19%3A11122841-11123341&hgt.suggest=&pix=800&Submit=submit&hgsid=184845147) | [SMARCA4](http://www.pdgene.org/view?gene=SMARCA4) | C | - | 10 | A vs. T | 0.05 (A) | - | >1 (-) | 0 (-) | <0.05 |
| [rs12609500](http://www.pdgene.org/view?poly=rs12609500) | [chr19:11173928](http://genome.ucsc.edu/cgi-bin/hgTracks?org=human&hgt.customText=http://www.pdgene.org/tracks?hg=hg19&db=hg19&position=chr19%3A11173678-11174178&hgt.suggest=&pix=800&Submit=submit&hgsid=184845147) | [SMARCA4](http://www.pdgene.org/view?gene=SMARCA4) | C | - | 12 | T vs. C | - | - | <1 (-) | 0 (-) | <0.05 |
| [rs73013198](http://www.pdgene.org/view?poly=rs73013198) | [chr19:11174742](http://genome.ucsc.edu/cgi-bin/hgTracks?org=human&hgt.customText=http://www.pdgene.org/tracks?hg=hg19&db=hg19&position=chr19%3A11174492-11174992&hgt.suggest=&pix=800&Submit=submit&hgsid=184845147) | [SMARCA4](http://www.pdgene.org/view?gene=SMARCA4) | C | - | 12 | T vs. C | 0.242 (T) | 0.05 (T) | <1 (-) | 0 (-) | <0.05 |
| [rs73013202](http://www.pdgene.org/view?poly=rs73013202) | [chr19:11179709](http://genome.ucsc.edu/cgi-bin/hgTracks?org=human&hgt.customText=http://www.pdgene.org/tracks?hg=hg19&db=hg19&position=chr19%3A11179459-11179959&hgt.suggest=&pix=800&Submit=submit&hgsid=184845147) | [SMARCA4[+3638bp]](http://www.pdgene.org/view?gene=SMARCA4) | C | - | 12 | C vs. G | 0.25 (G) | 0.067 (G) | >1 (-) | 0 (-) | <0.05 |
| [rs141989097](http://www.pdgene.org/view?poly=rs141989097) | [chr19:11069069](http://genome.ucsc.edu/cgi-bin/hgTracks?org=human&hgt.customText=http://www.pdgene.org/tracks?hg=hg19&db=hg19&position=chr19%3A11068819-11069319&hgt.suggest=&pix=800&Submit=submit&hgsid=184845147) | [SMARCA4[-2529bp]](http://www.pdgene.org/view?gene=SMARCA4) | C | - | 12 | T vs. C | - | - | >1 (-) | 23 (-) | <0.05 |
| [rs112186070](http://www.pdgene.org/view?poly=rs112186070) | [chr19:11168261](http://genome.ucsc.edu/cgi-bin/hgTracks?org=human&hgt.customText=http://www.pdgene.org/tracks?hg=hg19&db=hg19&position=chr19%3A11168011-11168511&hgt.suggest=&pix=800&Submit=submit&hgsid=184845147) | [SMARCA4](http://www.pdgene.org/view?gene=SMARCA4) | C | - | 12 | T vs. C | - | - | <1 (-) | 0 (-) | <0.05 |
| [rs113113862](http://www.pdgene.org/view?poly=rs113113862) | [chr19:11183577](http://genome.ucsc.edu/cgi-bin/hgTracks?org=human&hgt.customText=http://www.pdgene.org/tracks?hg=hg19&db=hg19&position=chr19%3A11183327-11183827&hgt.suggest=&pix=800&Submit=submit&hgsid=184845147) | [SMARCA4[+7506bp]](http://www.pdgene.org/view?gene=SMARCA4) | C | - | 12 | A vs. G | - | - | <1 (-) | 0 (-) | <0.05 |
| [rs73015007](http://www.pdgene.org/view?poly=rs73015007) | [chr19:11183837](http://genome.ucsc.edu/cgi-bin/hgTracks?org=human&hgt.customText=http://www.pdgene.org/tracks?hg=hg19&db=hg19&position=chr19%3A11183587-11184087&hgt.suggest=&pix=800&Submit=submit&hgsid=184845147) | [SMARCA4[+7766bp]](http://www.pdgene.org/view?gene=SMARCA4) | C | - | 12 | A vs. G | 0.25 (A) | - | <1 (-) | 0 (-) | <0.05 |
| [rs1122608](http://www.pdgene.org/view?poly=rs1122608) | [chr19:11163601](http://genome.ucsc.edu/cgi-bin/hgTracks?org=human&hgt.customText=http://www.pdgene.org/tracks?hg=hg19&db=hg19&position=chr19%3A11163351-11163851&hgt.suggest=&pix=800&Submit=submit&hgsid=184845147) | [SMARCA4](http://www.pdgene.org/view?gene=SMARCA4) | C | - | 12 | T vs. G | 0.25 (T) | 0.108 (T) | <1 (-) | 0 (-) | <0.05 |
| [rs3786728](http://www.pdgene.org/view?poly=rs3786728) | [chr19:11168038](http://genome.ucsc.edu/cgi-bin/hgTracks?org=human&hgt.customText=http://www.pdgene.org/tracks?hg=hg19&db=hg19&position=chr19%3A11167788-11168288&hgt.suggest=&pix=800&Submit=submit&hgsid=184845147) | [SMARCA4](http://www.pdgene.org/view?gene=SMARCA4) | C | - | 12 | A vs. G | 0.25 (G) | 0.067 (G) | >1 (-) | 0 (-) | <0.05 |
| [rs3786722](http://www.pdgene.org/view?poly=rs3786722) | [chr19:11161537](http://genome.ucsc.edu/cgi-bin/hgTracks?org=human&hgt.customText=http://www.pdgene.org/tracks?hg=hg19&db=hg19&position=chr19%3A11161287-11161787&hgt.suggest=&pix=800&Submit=submit&hgsid=184845147) | [SMARCA4](http://www.pdgene.org/view?gene=SMARCA4) | C | - | 12 | A vs. C | 0.25 (A) | 0.108 (A) | <1 (-) | 0 (-) | <0.05 |
| [rs112369586](http://www.pdgene.org/view?poly=rs112369586) | [chr19:11164460](http://genome.ucsc.edu/cgi-bin/hgTracks?org=human&hgt.customText=http://www.pdgene.org/tracks?hg=hg19&db=hg19&position=chr19%3A11164210-11164710&hgt.suggest=&pix=800&Submit=submit&hgsid=184845147) | [SMARCA4](http://www.pdgene.org/view?gene=SMARCA4) | C | - | 12 | A vs. G | 0.25 (A) | 0.092 (A) | <1 (-) | 0 (-) | <0.05 |
| [rs12609863](http://www.pdgene.org/view?poly=rs12609863) | [chr19:11167219](http://genome.ucsc.edu/cgi-bin/hgTracks?org=human&hgt.customText=http://www.pdgene.org/tracks?hg=hg19&db=hg19&position=chr19%3A11166969-11167469&hgt.suggest=&pix=800&Submit=submit&hgsid=184845147) | [SMARCA4](http://www.pdgene.org/view?gene=SMARCA4) | C | - | 12 | T vs. C | 0.258 (T) | 0.108 (T) | <1 (-) | 0 (-) | <0.05 |
| [rs117210556](http://www.pdgene.org/view?poly=rs117210556) | [chr19:11126197](http://genome.ucsc.edu/cgi-bin/hgTracks?org=human&hgt.customText=http://www.pdgene.org/tracks?hg=hg19&db=hg19&position=chr19%3A11125947-11126447&hgt.suggest=&pix=800&Submit=submit&hgsid=184845147) | [SMARCA4](http://www.pdgene.org/view?gene=SMARCA4) | C | - | 9 | T vs. C | 0.058 (C) | - | <1 (-) | 0 (-) | <0.05 |
| [rs55948246](http://www.pdgene.org/view?poly=rs55948246) | [chr19:11159889](http://genome.ucsc.edu/cgi-bin/hgTracks?org=human&hgt.customText=http://www.pdgene.org/tracks?hg=hg19&db=hg19&position=chr19%3A11159639-11160139&hgt.suggest=&pix=800&Submit=submit&hgsid=184845147) | [SMARCA4](http://www.pdgene.org/view?gene=SMARCA4) | C | - | 12 | T vs. C | 0.25 (C) | 0.092 (C) | >1 (-) | 0 (-) | <0.05 |
| [rs12052058](http://www.pdgene.org/view?poly=rs12052058) | [chr19:11159525](http://genome.ucsc.edu/cgi-bin/hgTracks?org=human&hgt.customText=http://www.pdgene.org/tracks?hg=hg19&db=hg19&position=chr19%3A11159275-11159775&hgt.suggest=&pix=800&Submit=submit&hgsid=184845147) | [SMARCA4](http://www.pdgene.org/view?gene=SMARCA4) | C | - | 12 | T vs. G | 0.25 (T) | 0.1 (T) | <1 (-) | 0 (-) | <0.05 |
| [rs12052201](http://www.pdgene.org/view?poly=rs12052201) | [chr19:11159096](http://genome.ucsc.edu/cgi-bin/hgTracks?org=human&hgt.customText=http://www.pdgene.org/tracks?hg=hg19&db=hg19&position=chr19%3A11158846-11159346&hgt.suggest=&pix=800&Submit=submit&hgsid=184845147) | [SMARCA4](http://www.pdgene.org/view?gene=SMARCA4) | C | - | 12 | T vs. G | 0.25 (T) | 0.1 (T) | <1 (-) | 0 (-) | <0.05 |
| [rs79227604](http://www.pdgene.org/view?poly=rs79227604) | [chr19:11065183](http://genome.ucsc.edu/cgi-bin/hgTracks?org=human&hgt.customText=http://www.pdgene.org/tracks?hg=hg19&db=hg19&position=chr19%3A11064933-11065433&hgt.suggest=&pix=800&Submit=submit&hgsid=184845147) | [SMARCA4[-6415bp]](http://www.pdgene.org/view?gene=SMARCA4) | C | - | 12 | T vs. C | 0.1 (T) | - | >1 (-) | 0 (-) | <0.05 |
| [rs12052200](http://www.pdgene.org/view?poly=rs12052200) | [chr19:11159076](http://genome.ucsc.edu/cgi-bin/hgTracks?org=human&hgt.customText=http://www.pdgene.org/tracks?hg=hg19&db=hg19&position=chr19%3A11158826-11159326&hgt.suggest=&pix=800&Submit=submit&hgsid=184845147) | [SMARCA4](http://www.pdgene.org/view?gene=SMARCA4) | C | - | 12 | A vs. G | 0.325 (A) | 0.308 (A) | <1 (-) | 0 (-) | <0.05 |
| [rs76614166](http://www.pdgene.org/view?poly=rs76614166) | [chr19:11104243](http://genome.ucsc.edu/cgi-bin/hgTracks?org=human&hgt.customText=http://www.pdgene.org/tracks?hg=hg19&db=hg19&position=chr19%3A11103993-11104493&hgt.suggest=&pix=800&Submit=submit&hgsid=184845147) | [SMARCA4](http://www.pdgene.org/view?gene=SMARCA4) | C | - | 11 | A vs. G | 0.05 (G) | - | <1 (-) | 3 (-) | <0.05 |
| [rs118158676](http://www.pdgene.org/view?poly=rs118158676) | [chr19:11079212](http://genome.ucsc.edu/cgi-bin/hgTracks?org=human&hgt.customText=http://www.pdgene.org/tracks?hg=hg19&db=hg19&position=chr19%3A11078962-11079462&hgt.suggest=&pix=800&Submit=submit&hgsid=184845147) | [SMARCA4](http://www.pdgene.org/view?gene=SMARCA4) | C | - | 11 | T vs. C | 0.025 (T) | - | <1 (-) | 0 (-) | <0.05 |
| [rs3786721](http://www.pdgene.org/view?poly=rs3786721) | [chr19:11146499](http://genome.ucsc.edu/cgi-bin/hgTracks?org=human&hgt.customText=http://www.pdgene.org/tracks?hg=hg19&db=hg19&position=chr19%3A11146249-11146749&hgt.suggest=&pix=800&Submit=submit&hgsid=184845147) | [SMARCA4](http://www.pdgene.org/view?gene=SMARCA4) | C | - | 12 | T vs. C | 0.45 (T) | 0.125 (T) | >1 (-) | 0 (-) | <0.05 |
| [rs10417443](http://www.pdgene.org/view?poly=rs10417443) | [chr19:11129429](http://genome.ucsc.edu/cgi-bin/hgTracks?org=human&hgt.customText=http://www.pdgene.org/tracks?hg=hg19&db=hg19&position=chr19%3A11129179-11129679&hgt.suggest=&pix=800&Submit=submit&hgsid=184845147) | [SMARCA4](http://www.pdgene.org/view?gene=SMARCA4) | C | - | 12 | C vs. G | 0.483 (C) | 0.117 (C) | >1 (-) | 0 (-) | <0.05 |
| [rs34285220](http://www.pdgene.org/view?poly=rs34285220) | [chr19:11184260](http://genome.ucsc.edu/cgi-bin/hgTracks?org=human&hgt.customText=http://www.pdgene.org/tracks?hg=hg19&db=hg19&position=chr19%3A11184010-11184510&hgt.suggest=&pix=800&Submit=submit&hgsid=184845147) | [SMARCA4[+8189bp]](http://www.pdgene.org/view?gene=SMARCA4) | C | - | 11 | T vs. C | - | - | >1 (-) | 9 (-) | <0.05 |
| [rs78301016](http://www.pdgene.org/view?poly=rs78301016) | [chr19:11063046](http://genome.ucsc.edu/cgi-bin/hgTracks?org=human&hgt.customText=http://www.pdgene.org/tracks?hg=hg19&db=hg19&position=chr19%3A11062796-11063296&hgt.suggest=&pix=800&Submit=submit&hgsid=184845147) | [SMARCA4[-8552bp]](http://www.pdgene.org/view?gene=SMARCA4) | C | - | 12 | A vs. G | 0.183 (A) | 0.025 (A) | >1 (-) | 30 (-) | <0.05 |

**Table S4. BLVRA PD risk SNPs**

| [**Polymorphism**](http://www.pdgene.org/view?gene=BLVRA) | [**Location (hg19)**](http://www.pdgene.org/view?gene=BLVRA) | **Gene** | [**Ethnicity**](http://www.pdgene.org/view?gene=BLVRA) | [**# Samples**](http://www.pdgene.org/view?gene=BLVRA) | [**# Studies**](http://www.pdgene.org/view?gene=BLVRA) | [**Allele contrast**](http://www.pdgene.org/view?gene=BLVRA) | **1000G CEU** | **1000G CHB+JPT** | **Meta OR (95%CI)** | **I2 (95%CI)** | [**Meta P-value**](http://www.pdgene.org/view?gene=BLVRA) |
| --- | --- | --- | --- | --- | --- | --- | --- | --- | --- | --- | --- |
| [rs1813599](http://www.pdgene.org/view?poly=rs1813599) | [chr7:43869788](http://genome.ucsc.edu/cgi-bin/hgTracks?org=human&hgt.customText=http://www.pdgene.org/tracks?hg=hg19&db=hg19&position=chr7%3A43869538-43870038&hgt.suggest=&pix=800&Submit=submit&hgsid=184845147) | [BLVRA[+22849bp]](http://www.pdgene.org/view?gene=BLVRA) | C | - | 13 | A vs. G | - | - | >1 (-) | 6 (-) | <0.05 |
| [rs10951743](http://www.pdgene.org/view?poly=rs10951743) | [chr7:43870113](http://genome.ucsc.edu/cgi-bin/hgTracks?org=human&hgt.customText=http://www.pdgene.org/tracks?hg=hg19&db=hg19&position=chr7%3A43869863-43870363&hgt.suggest=&pix=800&Submit=submit&hgsid=184845147) | [BLVRA[+23174bp]](http://www.pdgene.org/view?gene=BLVRA) | C | - | 13 | T vs. C | 0.383 (C) | - | <1 (-) | 0 (-) | <0.05 |
| [rs6978907](http://www.pdgene.org/view?poly=rs6978907) | [chr7:43835607](http://genome.ucsc.edu/cgi-bin/hgTracks?org=human&hgt.customText=http://www.pdgene.org/tracks?hg=hg19&db=hg19&position=chr7%3A43835357-43835857&hgt.suggest=&pix=800&Submit=submit&hgsid=184845147) | [BLVRA](http://www.pdgene.org/view?gene=BLVRA) | C | - | 13 | A vs. G | - | - | <1 (-) | 0 (-) | <0.05 |
| [rs146815437](http://www.pdgene.org/view?poly=rs146815437) | [chr7:43829380](http://genome.ucsc.edu/cgi-bin/hgTracks?org=human&hgt.customText=http://www.pdgene.org/tracks?hg=hg19&db=hg19&position=chr7%3A43829130-43829630&hgt.suggest=&pix=800&Submit=submit&hgsid=184845147) | [BLVRA](http://www.pdgene.org/view?gene=BLVRA) | C | - | 10 | A vs. C | - | - | <1 (-) | 43 (-) | <0.05 |
| [rs6945433](http://www.pdgene.org/view?poly=rs6945433) | [chr7:43858926](http://genome.ucsc.edu/cgi-bin/hgTracks?org=human&hgt.customText=http://www.pdgene.org/tracks?hg=hg19&db=hg19&position=chr7%3A43858676-43859176&hgt.suggest=&pix=800&Submit=submit&hgsid=184845147) | [BLVRA[+11987bp]](http://www.pdgene.org/view?gene=BLVRA) | C | - | 13 | T vs. C | 0.333 (T) | 0.325 (T) | >1 (-) | 0 (-) | <0.05 |
| [rs143530356](http://www.pdgene.org/view?poly=rs143530356) | [chr7:43829579](http://genome.ucsc.edu/cgi-bin/hgTracks?org=human&hgt.customText=http://www.pdgene.org/tracks?hg=hg19&db=hg19&position=chr7%3A43829329-43829829&hgt.suggest=&pix=800&Submit=submit&hgsid=184845147) | [BLVRA](http://www.pdgene.org/view?gene=BLVRA) | C | - | 4 | C vs. G | - | - | >1 (-) | 0 (-) | <0.05 |
| [rs3094952](http://www.pdgene.org/view?poly=rs3094952) | [chr7:43837562](http://genome.ucsc.edu/cgi-bin/hgTracks?org=human&hgt.customText=http://www.pdgene.org/tracks?hg=hg19&db=hg19&position=chr7%3A43837312-43837812&hgt.suggest=&pix=800&Submit=submit&hgsid=184845147) | [BLVRA](http://www.pdgene.org/view?gene=BLVRA) | C | - | 13 | A vs. G | - | - | <1 (-) | 0 (-) | <0.05 |
| [rs34865291](http://www.pdgene.org/view?poly=rs34865291) | [chr7:43864079](http://genome.ucsc.edu/cgi-bin/hgTracks?org=human&hgt.customText=http://www.pdgene.org/tracks?hg=hg19&db=hg19&position=chr7%3A43863829-43864329&hgt.suggest=&pix=800&Submit=submit&hgsid=184845147) | [BLVRA[+17140bp]](http://www.pdgene.org/view?gene=BLVRA) | C | - | 13 | A vs. G | 0.025 (G) | - | <1 (-) | 0 (-) | <0.05 |
| [rs147976254](http://www.pdgene.org/view?poly=rs147976254) | [chr7:43866701](http://genome.ucsc.edu/cgi-bin/hgTracks?org=human&hgt.customText=http://www.pdgene.org/tracks?hg=hg19&db=hg19&position=chr7%3A43866451-43866951&hgt.suggest=&pix=800&Submit=submit&hgsid=184845147) | [BLVRA[+19762bp]](http://www.pdgene.org/view?gene=BLVRA) | C | - | 12 | A vs. G | - | - | >1 (-) | 0 (-) | <0.05 |
| [rs2282922](http://www.pdgene.org/view?poly=rs2282922) | [chr7:43844467](http://genome.ucsc.edu/cgi-bin/hgTracks?org=human&hgt.customText=http://www.pdgene.org/tracks?hg=hg19&db=hg19&position=chr7%3A43844217-43844717&hgt.suggest=&pix=800&Submit=submit&hgsid=184845147) | [BLVRA](http://www.pdgene.org/view?gene=BLVRA) | C | - | 13 | T vs. C | 0.283 (C) | 0.267 (C) | >1 (-) | 0 (-) | <0.05 |
| [rs1181602](http://www.pdgene.org/view?poly=rs1181602) | [chr7:43795677](http://genome.ucsc.edu/cgi-bin/hgTracks?org=human&hgt.customText=http://www.pdgene.org/tracks?hg=hg19&db=hg19&position=chr7%3A43795427-43795927&hgt.suggest=&pix=800&Submit=submit&hgsid=184845147) | [BLVRA[-2602bp]](http://www.pdgene.org/view?gene=BLVRA) | C | - | 13 | T vs. G | - | - | <1 (-) | 0 (-) | <0.05 |
| [rs2730625](http://www.pdgene.org/view?poly=rs2730625) | [chr7:43845944](http://genome.ucsc.edu/cgi-bin/hgTracks?org=human&hgt.customText=http://www.pdgene.org/tracks?hg=hg19&db=hg19&position=chr7%3A43845694-43846194&hgt.suggest=&pix=800&Submit=submit&hgsid=184845147) | [BLVRA](http://www.pdgene.org/view?gene=BLVRA) | C | - | 13 | T vs. C | 0.283 (C) | 0.267 (C) | >1 (-) | 0 (-) | <0.05 |
| [rs2299149](http://www.pdgene.org/view?poly=rs2299149) | [chr7:43845185](http://genome.ucsc.edu/cgi-bin/hgTracks?org=human&hgt.customText=http://www.pdgene.org/tracks?hg=hg19&db=hg19&position=chr7%3A43844935-43845435&hgt.suggest=&pix=800&Submit=submit&hgsid=184845147) | [BLVRA](http://www.pdgene.org/view?gene=BLVRA) | C | - | 13 | A vs. G | 0.283 (G) | 0.267 (G) | >1 (-) | 0 (-) | <0.05 |
| [rs849162](http://www.pdgene.org/view?poly=rs849162) | [chr7:43821852](http://genome.ucsc.edu/cgi-bin/hgTracks?org=human&hgt.customText=http://www.pdgene.org/tracks?hg=hg19&db=hg19&position=chr7%3A43821602-43822102&hgt.suggest=&pix=800&Submit=submit&hgsid=184845147) | [BLVRA](http://www.pdgene.org/view?gene=BLVRA) | C | - | 13 | A vs. G | 0.283 (A) | 0.267 (A) | <1 (-) | 0 (-) | <0.05 |
| [rs699512](http://www.pdgene.org/view?poly=rs699512) | [chr7:43810764](http://genome.ucsc.edu/cgi-bin/hgTracks?org=human&hgt.customText=http://www.pdgene.org/tracks?hg=hg19&db=hg19&position=chr7%3A43810514-43811014&hgt.suggest=&pix=800&Submit=submit&hgsid=184845147) | [BLVRA](http://www.pdgene.org/view?gene=BLVRA) | C | - | 13 | A vs. G | 0.283 (G) | 0.267 (G) | >1 (-) | 0 (-) | <0.05 |
| [rs699510](http://www.pdgene.org/view?poly=rs699510) | [chr7:43810270](http://genome.ucsc.edu/cgi-bin/hgTracks?org=human&hgt.customText=http://www.pdgene.org/tracks?hg=hg19&db=hg19&position=chr7%3A43810020-43810520&hgt.suggest=&pix=800&Submit=submit&hgsid=184845147) | [BLVRA](http://www.pdgene.org/view?gene=BLVRA) | C | - | 13 | T vs. C | 0.283 (C) | 0.267 (C) | >1 (-) | 0 (-) | <0.05 |
| [rs2528369](http://www.pdgene.org/view?poly=rs2528369) | [chr7:43848123](http://genome.ucsc.edu/cgi-bin/hgTracks?org=human&hgt.customText=http://www.pdgene.org/tracks?hg=hg19&db=hg19&position=chr7%3A43847873-43848373&hgt.suggest=&pix=800&Submit=submit&hgsid=184845147) | [BLVRA[+1184bp]](http://www.pdgene.org/view?gene=BLVRA) | C | - | 13 | T vs. C | 0.267 (T) | 0.267 (T) | <1 (-) | 0 (-) | <0.05 |
| [rs2246171](http://www.pdgene.org/view?poly=rs2246171) | [chr7:43840347](http://genome.ucsc.edu/cgi-bin/hgTracks?org=human&hgt.customText=http://www.pdgene.org/tracks?hg=hg19&db=hg19&position=chr7%3A43840097-43840597&hgt.suggest=&pix=800&Submit=submit&hgsid=184845147) | [BLVRA](http://www.pdgene.org/view?gene=BLVRA) | C | - | 13 | A vs. G | 0.267 (G) | 0.267 (G) | >1 (-) | 0 (-) | <0.05 |
| [rs3094951](http://www.pdgene.org/view?poly=rs3094951) | [chr7:43840777](http://genome.ucsc.edu/cgi-bin/hgTracks?org=human&hgt.customText=http://www.pdgene.org/tracks?hg=hg19&db=hg19&position=chr7%3A43840527-43841027&hgt.suggest=&pix=800&Submit=submit&hgsid=184845147) | [BLVRA](http://www.pdgene.org/view?gene=BLVRA) | C | - | 13 | T vs. G | 0.267 (G) | 0.267 (G) | >1 (-) | 0 (-) | <0.05 |
| [rs1181573](http://www.pdgene.org/view?poly=rs1181573) | [chr7:43803607](http://genome.ucsc.edu/cgi-bin/hgTracks?org=human&hgt.customText=http://www.pdgene.org/tracks?hg=hg19&db=hg19&position=chr7%3A43803357-43803857&hgt.suggest=&pix=800&Submit=submit&hgsid=184845147) | [BLVRA](http://www.pdgene.org/view?gene=BLVRA) | C | - | 13 | A vs. G | 0.283 (G) | 0.267 (G) | >1 (-) | 0 (-) | <0.05 |
| [rs1317916](http://www.pdgene.org/view?poly=rs1317916) | [chr7:43839511](http://genome.ucsc.edu/cgi-bin/hgTracks?org=human&hgt.customText=http://www.pdgene.org/tracks?hg=hg19&db=hg19&position=chr7%3A43839261-43839761&hgt.suggest=&pix=800&Submit=submit&hgsid=184845147) | [BLVRA](http://www.pdgene.org/view?gene=BLVRA) | C | - | 13 | A vs. G | 0.267 (A) | 0.267 (A) | <1 (-) | 0 (-) | <0.05 |
| [rs1306743](http://www.pdgene.org/view?poly=rs1306743) | [chr7:43835218](http://genome.ucsc.edu/cgi-bin/hgTracks?org=human&hgt.customText=http://www.pdgene.org/tracks?hg=hg19&db=hg19&position=chr7%3A43834968-43835468&hgt.suggest=&pix=800&Submit=submit&hgsid=184845147) | [BLVRA](http://www.pdgene.org/view?gene=BLVRA) | C | - | 13 | A vs. T | 0.267 (T) | 0.267 (T) | >1 (-) | 0 (-) | <0.05 |
| [rs3107889](http://www.pdgene.org/view?poly=rs3107889) | [chr7:43838468](http://genome.ucsc.edu/cgi-bin/hgTracks?org=human&hgt.customText=http://www.pdgene.org/tracks?hg=hg19&db=hg19&position=chr7%3A43838218-43838718&hgt.suggest=&pix=800&Submit=submit&hgsid=184845147) | [BLVRA](http://www.pdgene.org/view?gene=BLVRA) | C | - | 13 | A vs. C | 0.275 (A) | 0.267 (A) | <1 (-) | 0 (-) | <0.05 |
| [rs1306742](http://www.pdgene.org/view?poly=rs1306742) | [chr7:43833162](http://genome.ucsc.edu/cgi-bin/hgTracks?org=human&hgt.customText=http://www.pdgene.org/tracks?hg=hg19&db=hg19&position=chr7%3A43832912-43833412&hgt.suggest=&pix=800&Submit=submit&hgsid=184845147) | [BLVRA](http://www.pdgene.org/view?gene=BLVRA) | C | - | 13 | T vs. C | 0.267 (T) | 0.267 (T) | <1 (-) | 0 (-) | <0.05 |
| [rs2730604](http://www.pdgene.org/view?poly=rs2730604) | [chr7:43850459](http://genome.ucsc.edu/cgi-bin/hgTracks?org=human&hgt.customText=http://www.pdgene.org/tracks?hg=hg19&db=hg19&position=chr7%3A43850209-43850709&hgt.suggest=&pix=800&Submit=submit&hgsid=184845147) | [BLVRA[+3520bp]](http://www.pdgene.org/view?gene=BLVRA) | C | - | 13 | A vs. C | 0.292 (C) | 0.283 (C) | >1 (-) | 0 (-) | <0.05 |
| [rs1637530](http://www.pdgene.org/view?poly=rs1637530) | [chr7:43797453](http://genome.ucsc.edu/cgi-bin/hgTracks?org=human&hgt.customText=http://www.pdgene.org/tracks?hg=hg19&db=hg19&position=chr7%3A43797203-43797703&hgt.suggest=&pix=800&Submit=submit&hgsid=184845147) | [BLVRA[-826bp]](http://www.pdgene.org/view?gene=BLVRA) | C | - | 13 | T vs. C | 0.283 (T) | 0.208 (T) | <1 (-) | 0 (-) | <0.05 |
| [rs3107888](http://www.pdgene.org/view?poly=rs3107888) | [chr7:43842117](http://genome.ucsc.edu/cgi-bin/hgTracks?org=human&hgt.customText=http://www.pdgene.org/tracks?hg=hg19&db=hg19&position=chr7%3A43841867-43842367&hgt.suggest=&pix=800&Submit=submit&hgsid=184845147) | [BLVRA](http://www.pdgene.org/view?gene=BLVRA) | C | - | 13 | A vs. T | 0.292 (T) | 0.267 (T) | >1 (-) | 0 (-) | <0.05 |
| [rs598042](http://www.pdgene.org/view?poly=rs598042) | [chr7:43873740](http://genome.ucsc.edu/cgi-bin/hgTracks?org=human&hgt.customText=http://www.pdgene.org/tracks?hg=hg19&db=hg19&position=chr7%3A43873490-43873990&hgt.suggest=&pix=800&Submit=submit&hgsid=184845147) | [BLVRA[+26801bp]](http://www.pdgene.org/view?gene=BLVRA) | C | - | 12 | A vs. C | 0.375 (C) | 0.417 (C) | >1 (-) | 0 (-) | <0.05 |
| [rs627932](http://www.pdgene.org/view?poly=rs627932) | [chr7:43871686](http://genome.ucsc.edu/cgi-bin/hgTracks?org=human&hgt.customText=http://www.pdgene.org/tracks?hg=hg19&db=hg19&position=chr7%3A43871436-43871936&hgt.suggest=&pix=800&Submit=submit&hgsid=184845147) | [BLVRA[+24747bp]](http://www.pdgene.org/view?gene=BLVRA) | C | - | 13 | A vs. C | 0.375 (C) | 0.417 (C) | >1 (-) | 0 (-) | <0.05 |
| [rs1181598](http://www.pdgene.org/view?poly=rs1181598) | [chr7:43789513](http://genome.ucsc.edu/cgi-bin/hgTracks?org=human&hgt.customText=http://www.pdgene.org/tracks?hg=hg19&db=hg19&position=chr7%3A43789263-43789763&hgt.suggest=&pix=800&Submit=submit&hgsid=184845147) | [BLVRA[-8766bp]](http://www.pdgene.org/view?gene=BLVRA) | C | - | 13 | C vs. G | 0.275 (C) | 0.267 (C) | <1 (-) | 0 (-) | <0.05 |
| [rs2529584](http://www.pdgene.org/view?poly=rs2529584) | [chr7:43790304](http://genome.ucsc.edu/cgi-bin/hgTracks?org=human&hgt.customText=http://www.pdgene.org/tracks?hg=hg19&db=hg19&position=chr7%3A43790054-43790554&hgt.suggest=&pix=800&Submit=submit&hgsid=184845147) | [BLVRA[-7975bp]](http://www.pdgene.org/view?gene=BLVRA) | C | - | 13 | A vs. G | 0.3 (A) | 0.267 (A) | <1 (-) | 0 (-) | <0.05 |
| [rs1181600](http://www.pdgene.org/view?poly=rs1181600) | [chr7:43789915](http://genome.ucsc.edu/cgi-bin/hgTracks?org=human&hgt.customText=http://www.pdgene.org/tracks?hg=hg19&db=hg19&position=chr7%3A43789665-43790165&hgt.suggest=&pix=800&Submit=submit&hgsid=184845147) | [BLVRA[-8364bp]](http://www.pdgene.org/view?gene=BLVRA) | C | - | 13 | T vs. C | 0.275 (T) | 0.267 (T) | <1 (-) | 0 (-) | <0.05 |
| [rs648048](http://www.pdgene.org/view?poly=rs648048) | [chr7:43861861](http://genome.ucsc.edu/cgi-bin/hgTracks?org=human&hgt.customText=http://www.pdgene.org/tracks?hg=hg19&db=hg19&position=chr7%3A43861611-43862111&hgt.suggest=&pix=800&Submit=submit&hgsid=184845147) | [BLVRA[+14922bp]](http://www.pdgene.org/view?gene=BLVRA) | C | - | 13 | A vs. G | - | - | <1 (-) | 0 (-) | <0.05 |
| [rs623108](http://www.pdgene.org/view?poly=rs623108) | [chr7:43864699](http://genome.ucsc.edu/cgi-bin/hgTracks?org=human&hgt.customText=http://www.pdgene.org/tracks?hg=hg19&db=hg19&position=chr7%3A43864449-43864949&hgt.suggest=&pix=800&Submit=submit&hgsid=184845147) | [BLVRA[+17760bp]](http://www.pdgene.org/view?gene=BLVRA) | C | - | 13 | A vs. G | 0.375 (A) | 0.417 (A) | <1 (-) | 0 (-) | <0.05 |
| [rs730585](http://www.pdgene.org/view?poly=rs730585) | [chr7:43789018](http://genome.ucsc.edu/cgi-bin/hgTracks?org=human&hgt.customText=http://www.pdgene.org/tracks?hg=hg19&db=hg19&position=chr7%3A43788768-43789268&hgt.suggest=&pix=800&Submit=submit&hgsid=184845147) | [BLVRA[-9261bp]](http://www.pdgene.org/view?gene=BLVRA) | C | - | 13 | A vs. G | 0.283 (A) | 0.267 (A) | <1 (-) | 0 (-) | <0.05 |
| [rs620833](http://www.pdgene.org/view?poly=rs620833) | [chr7:43865233](http://genome.ucsc.edu/cgi-bin/hgTracks?org=human&hgt.customText=http://www.pdgene.org/tracks?hg=hg19&db=hg19&position=chr7%3A43864983-43865483&hgt.suggest=&pix=800&Submit=submit&hgsid=184845147) | [BLVRA[+18294bp]](http://www.pdgene.org/view?gene=BLVRA) | C | - | 13 | C vs. G | 0.375 (G) | 0.417 (G) | >1 (-) | 0 (-) | <0.05 |
| [rs609979](http://www.pdgene.org/view?poly=rs609979) | [chr7:43873390](http://genome.ucsc.edu/cgi-bin/hgTracks?org=human&hgt.customText=http://www.pdgene.org/tracks?hg=hg19&db=hg19&position=chr7%3A43873140-43873640&hgt.suggest=&pix=800&Submit=submit&hgsid=184845147) | [BLVRA[+26451bp]](http://www.pdgene.org/view?gene=BLVRA) | C | - | 13 | A vs. C | 0.375 (A) | 0.417 (A) | <1 (-) | 0 (-) | <0.05 |
| [rs673402](http://www.pdgene.org/view?poly=rs673402) | [chr7:43873157](http://genome.ucsc.edu/cgi-bin/hgTracks?org=human&hgt.customText=http://www.pdgene.org/tracks?hg=hg19&db=hg19&position=chr7%3A43872907-43873407&hgt.suggest=&pix=800&Submit=submit&hgsid=184845147) | [BLVRA[+26218bp]](http://www.pdgene.org/view?gene=BLVRA) | C | - | 13 | A vs. G | - | - | <1 (-) | 0 (-) | <0.05 |
| [rs849180](http://www.pdgene.org/view?poly=rs849180) | [chr7:43785288](http://genome.ucsc.edu/cgi-bin/hgTracks?org=human&hgt.customText=http://www.pdgene.org/tracks?hg=hg19&db=hg19&position=chr7%3A43785038-43785538&hgt.suggest=&pix=800&Submit=submit&hgsid=184845147) | [BLVRA[-12991bp]](http://www.pdgene.org/view?gene=BLVRA) | C | - | 13 | T vs. C | - | - | >1 (-) | 0 (-) | <0.05 |
| [rs849179](http://www.pdgene.org/view?poly=rs849179) | [chr7:43785019](http://genome.ucsc.edu/cgi-bin/hgTracks?org=human&hgt.customText=http://www.pdgene.org/tracks?hg=hg19&db=hg19&position=chr7%3A43784769-43785269&hgt.suggest=&pix=800&Submit=submit&hgsid=184845147) | [BLVRA[-13260bp]](http://www.pdgene.org/view?gene=BLVRA) | C | - | 13 | A vs. T | 0.275 (T) | 0.267 (T) | >1 (-) | 0 (-) | <0.05 |
| [rs1181535](http://www.pdgene.org/view?poly=rs1181535) | [chr7:43874751](http://genome.ucsc.edu/cgi-bin/hgTracks?org=human&hgt.customText=http://www.pdgene.org/tracks?hg=hg19&db=hg19&position=chr7%3A43874501-43875001&hgt.suggest=&pix=800&Submit=submit&hgsid=184845147) | [BLVRA[+27812bp]](http://www.pdgene.org/view?gene=BLVRA) | C | - | 13 | A vs. G | 0.375 (G) | 0.417 (G) | >1 (-) | 0 (-) | <0.05 |
| [rs1181550](http://www.pdgene.org/view?poly=rs1181550) | [chr7:43861325](http://genome.ucsc.edu/cgi-bin/hgTracks?org=human&hgt.customText=http://www.pdgene.org/tracks?hg=hg19&db=hg19&position=chr7%3A43861075-43861575&hgt.suggest=&pix=800&Submit=submit&hgsid=184845147) | [BLVRA[+14386bp]](http://www.pdgene.org/view?gene=BLVRA) | C | - | 13 | A vs. G | 0.375 (G) | 0.417 (G) | >1 (-) | 0 (-) | <0.05 |
| [rs1181596](http://www.pdgene.org/view?poly=rs1181596) | [chr7:43785376](http://genome.ucsc.edu/cgi-bin/hgTracks?org=human&hgt.customText=http://www.pdgene.org/tracks?hg=hg19&db=hg19&position=chr7%3A43785126-43785626&hgt.suggest=&pix=800&Submit=submit&hgsid=184845147) | [BLVRA[-12903bp]](http://www.pdgene.org/view?gene=BLVRA) | C | - | 13 | A vs. G | - | - | <1 (-) | 0 (-) | <0.05 |
| [rs1181534](http://www.pdgene.org/view?poly=rs1181534) | [chr7:43875597](http://genome.ucsc.edu/cgi-bin/hgTracks?org=human&hgt.customText=http://www.pdgene.org/tracks?hg=hg19&db=hg19&position=chr7%3A43875347-43875847&hgt.suggest=&pix=800&Submit=submit&hgsid=184845147) | [BLVRA[+28658bp]](http://www.pdgene.org/view?gene=BLVRA) | C | - | 13 | A vs. G | 0.383 (G) | 0.417 (G) | >1 (-) | 0 (-) | <0.05 |

**Table S5. Oligos used for the experiments**

| **gene** | **Forward Primer (5’→ 3’)** | **Reverse Primer (5’→ 3’)** |
| --- | --- | --- |
| αTUB | CGTTTGTCAAGCCTCATAGC | ACACCAGCCTGACCAACAT |
| BVR | ATAACGCGCTGGACATCCTC | TGCTATTATTGGAGGAACCGGC |
| αSYN | CCACAGTGGCTGAGAAGACC | AATTCCTTCCTGTGGGGCTC |
