## APPENDIX for "Attenuation of epigenetic regulator SMARCA4 and ERK-ETS signaling suppresses aging-related dopaminergic degeneration"

**APPENDIX: Fly genotype list**

**Figure 2**

Figure 2d, e

TH-Gal4/+

TH-Gal4; UAS-LucRNAi

TH-Gal4; UAS-Brm(wt)

TH-Gal4; UAS-Brm RNAi

TH-Gal4; UAS-Brm^DN^

TH-Gal4; UAS-Bvr II

TH-Gal4; UAS-Bvr III

TH-Gal4; UAS-Bvr RNAi

TH-Gal4; UAS-Bvr II; UAS-Brm^DN^

Figure 2f:

TH-Gal4; UAS-RFP/Brm::GFP

TH-Gal4, UAS-αSynA30P; UAS-RFP/Brm::GFP

TH-Gal4, UAS-RFP; UAS-dLrrk2^I1915T^/Brm::GFP

*pink1*^B9^; TH-Gal4, UAS-RFP; Brm::GFP

TH-Gal4, UAS-Parkin RNAi; UAS-RFP/Brm::GFP

**Figure 3**

Figure 3a:

TH-Gal4, UAS-αSynA30P/+

TH-Gal4, UAS-αSynA30P; UAS-lucRNAi

TH-Gal4, UAS-αSynA30P; UAS-Brm(wt)

TH-Gal4, UAS-αSynA30P; UAS-Brm RNAi

TH-Gal4, UAS-αSynA30P; UAS-Brm^DN^

Figure 3b:

TH-Gal4; UAS-dLrrk2^I1915T^ /+

TH-Gal4; UAS-dLrrk2^I1915T^/UAS-lucRNAi

TH-Gal4; UAS-dLrrk2^I1915T^/UAS-Brm(wt)

TH-Gal4; UAS- dLrrk2^I1915T^ / UAS-Brm RNAi

TH-Gal4, UAS-dLrrk2^I1915T^; UAS-Brm^DN^

Figure 3c:

TH-Gal4, UAS-Parkin RNAi /+

TH-Gal4, UAS-Parkin RNAi; UAS-lucRNAi

TH-Gal4, UAS-Parkin RNAi; UAS-Brm(wt)

TH-Gal4, UAS-Parkin RNAi; UAS-Brm RNAi

TH-Gal4, UAS- Parkin RNAi; UAS-Brm^DN^

Figure 3d:

*pink1*^B9^; TH-Gal4/+

*pink1*^B9^; TH-Gal4/UAS-lucRNAi

*pink1*^B9^; TH-Gal4/UAS-Brm(wt)

*pink1*^B9^; TH-Gal4/UAS-Brm RNAi

*pink1*^B9^; TH-Gal4/ UAS-Brm^DN^

Figure 3e:

TH-Gal4, UAS-αSynA30P/+

TH-Gal4, UAS-αSynA30P/UAS-dBVR RNAi

TH-Gal4, UAS-αSynA30P/UAS-dBVR (III)

TH-Gal4, UAS-αSynA30P/UAS-dBVR (II)

TH-Gal4, UAS-αSynA30P/UAS-dBVR (II); UAS-Brm^DN^

TH-Gal4, UAS-αSynA30P/UAS-dBVR (II); UAS-MEK RNAi

TH-Gal4, UAS-αSynA30P/UAS-dBVR (III); UAS-Aop^wt^

Figure 3f:

TH-Gal4; UAS- UAS-dLrrk2^I1915T^ /+

TH-Gal4; UAS-dLrrk2^I1915T^/UAS-dBVR RNAi

TH-Gal4; UAS-dLrrk2^I1915T^/UAS-dBVR(III)

TH-Gal4; UAS-dLrrk2^I1915T^ /UAS-dBVR(II)

TH-Gal4; UAS-dLrrk2^I1915T^ / UAS-dBVR (II); UAS-Brm^DN^

TH-Gal4; UAS-dLrrk2^I1915T^/UAS-dBVR(II); UAS-MEK RNAi

TH-Gal4; UAS-dLrrk2^I1915T^/UAS-dBVR(III); UAS-Aop^wt^

Figure 3g:

TH-Gal4, UAS-Parkin RNAi/+

TH-Gal4, UAS-Parkin RNAi/UAS-LucRNAi

TH-Gal4; UAS-Parkin RNAi/UAS-dBVR RNAi

TH-Gal4; UAS-Parkin RNAi /UAS-dBVR(III)

TH-Gal4; UAS-Parkin RNAi /UAS-dBVR(II)

TH-Gal4; UAS-Parkin RNAi / UAS-dBVR (II); UAS-Brm^DN^

TH-Gal4, UAS-Parkin RNAi/UAS-dBVR(II); UAS-MEK RNAi

TH-Gal4, UAS-Parkin RNAi/UAS-dBVR(III); UAS-Aop^wt^

Figure 3h:

*pink1*^B9^; TH-Gal4/+

*pink1*^B9^; TH-Gal4/UAS-lucRNAi

*pink1*^B9^; TH-Gal4/UAS-dBVR RNAi

*pink1*^B9^; TH-Gal4/UAS-dBVR(III)

*pink1*^B9^; TH-Gal4/UAS-dBVR(II)

*pink1*^B9^; TH-Gal4/UAS-dBVR(II); UAS-Brm^DN^

*pink1*^B9^; TH-Gal4/UAS-dBVR(II); UAS-MEK RNAi

*pink1*^B9^; TH-Gal4/UAS-dBVR(II); UAS-Aop^wt^

Figure 3i, j:

TH-Gal4, UAS-RFP; Pnt::EGFP

TH-Gal4, UAS-RFP; Pnt::EGFP/UAS-dBrm RNAi

Figure 3k, l:

TH-Gal4, UAS-RFP; Pnt::EGFP

TH-Gal4, UAS-RFP; Pnt::EGFP/UAS-dBrm (wt)

Figure 3m-n:

TH-Gal4, UAS-RFP; Pnt::EGFP

TH-Gal4, UAS-RFP; Pnt::EGFP/UAS-dBvr RNAi

TH-Gal4, UAS-RFP; Pnt::EGFP/UAS-dBvr II

**Figure 4**

Figure 4a, b:

*elav*-Gal4/+

*elav*-Gal4/UAS-lucRNAi

*elav*-Gal4/UAS-αSynA30P

*elav*-Gal4/UAS-dLrrk2^I1915T^

*elav*-Gal4/UAS-dPINK1 RNAi

*elav*-Gal4; UAS-dParkin RNAi

Figure 4c-e:

TH-Gal4; UAS-RFP/Pnt::EGFP

TH-Gal4, UAS-αSynA30P; UAS-RFP/Pnt::EGFP

TH-Gal4, UAS-RFP; UAS-dLrrk2^I1915T^/Pnt::EGFP

*pink1*^B9^; TH-Gal4, UAS-RFP; Pnt::EGFP

TH-Gal4, UAS-Parkin RNAi; UAS-RFP/Pnt::EGFP

Figure 4f:

TH-Gal4, UAS-αSynA30P/+

TH-Gal4, UAS-αSynA30P; UAS-lucRNAi

TH-Gal4, UAS-αSynA30P; UAS-Erk RNAi

TH-Gal4, UAS-αSynA30P; UAS-MEK RNAi

Figure 4g:

TH-Gal4; UAS- UAS-dLrrk2^I1915T^ /+

TH-Gal4; UAS- UAS-dLrrk2^I1915T^/UAS-lucRNAi

TH-Gal4; UAS- UAS-dLrrk2^I1915T^/UAS-Erk RNAi

TH-Gal4; UAS- UAS-dLrrk2^I1915T^/ UAS-MEK RNAi

Figure 4h:

TH-Gal4, UAS-Parkin RNAi /+

TH-Gal4, UAS-Parkin RNAi; UAS-lucRNAi

TH-Gal4, UAS-Parkin RNAi; UAS-Erk RNAi

TH-Gal4, UAS-Parkin RNAi; UAS-MEK RNAi

Figure 4i:

*pink1*^B9^; TH-Gal4/+

*pink1*^B9^; TH-Gal4/UAS-lucRNAi

*pink1*^B9^; TH-Gal4/UAS-Erk RNAi

*pink1*^B9^; TH-Gal4/UAS-MEK RNAi

Figure 4j:

TH-Gal4, UAS-αSynA30P/+

TH-Gal4, UAS-αSynA30P/UAS-LucRNAi

TH-Gal4, UAS-αSynA30P/UAS-Pnt RNAi

TH-Gal4, UAS-αSynA30P/UAS-Aop RNAi

TH-Gal4, UAS-αSynA30P/UAS-Aop^wt^

Figure 4k:

TH-Gal4; UAS-dLrrk2^I1915T^ /+

TH-Gal4; UAS-dLrrk2^I1915T^ /LucRNAi

TH-Gal4; UAS-dLrrk2^I1915T^/UAS-Pnt RNAi

TH-Gal4; UAS-dLrrk2^I1915T^/UAS-Aop RNAi

TH-Gal4; UAS-dLrrk2^I1915T^ /UAS-Aop^wt^

Figure 4l:

TH-Gal4, UAS-Parkin RNAi/+

TH-Gal4, UAS-Parkin RNAi/UAS-LucRNAi

TH-Gal4; UAS-Parkin RNAi/UAS-Pnt RNAi

TH-Gal4; UAS-Parkin RNAi /UAS-Aop RNAi

TH-Gal4, UAS-Parkin RNAi/UAS-Aop^wt^

Figure 4m:

*pink1*^B9^; TH-Gal4/+

*pink1*^B9^; TH-Gal4/UAS-lucRNAi

*pink1*^B9^; TH-Gal4/UAS-Pnt RNAi

*pink1*^B9^; TH-Gal4/UAS-Aop RNAi

*pink1*^B9^; TH-Gal4/UAS-Aop^wt^

Figure 4n-p:

TH-Gal4/+

TH-Gal4; UAS-αSynA30P

TH-Gal4; UAS-dLrrk2^I1915T^

TH-Gal4; UAS-dParkin RNAi

*pink1*^B9^; TH-Gal4/+

**Figure S5**

TH-Gal4/+

TH-Gal4; UAS-αSynA30P

TH-Gal4; UAS-dLrrk2^I1915T^

TH-Gal4; UAS-dParkin RNAi

TH-Gal4; UAS-dPINK1 RNAi

*pink1*^B9^; TH-Gal4/+

**Figure S6**

TH-Gal4; UAS-RFP/Tub-mito-roGFP2

TH-Gal4, UAS-αSynA30P; UAS-RFP/ Tub-mito-roGFP2

TH-Gal4, UAS-RFP; UAS-dLrrk2^I1915T^/ Tub-mito-roGFP2

TH-Gal4, UAS-PINK1 RNAi; UAS-RFP/ Tub-mito-roGFP2

TH-Gal4, UAS-Parkin RNAi; UAS-RFP/ Tub-mito-roGFP2

**Figure S7**

TH-Gal4; UAS-RFP/UAS-roGFP2

TH-Gal4, UAS-αSynA30P; UAS-RFP/ UAS-roGFP2

TH-Gal4, UAS-RFP; UAS-dLrrk2^I1915T^/ UAS-roGFP2

TH-Gal4, UAS-PINK1 RNAi; UAS-RFP/ UAS-roGFP2

TH-Gal4, UAS-Parkin RNAi; UAS-RFP/ UAS-roGFP2

**Figure S8**

TH-Gal4; UAS-RFP/GstD-GFP

TH-Gal4, UAS-αSynA30P; UAS-RFP/GstD-GFP

TH-Gal4, UAS-RFP; UAS-dLrrk2^I1915T^/GstD-GFP

TH-Gal4, UAS-PINK1 RNAi; UAS-RFP/GstD-GFP

TH-Gal4, UAS-Parkin RNAi; UAS-RFP/GstD-GFP
